# Structural, biophysical and biological analysis and characterisation of IRF4 DNA-binding domain mutations associated with multiple myeloma

**DOI:** 10.1101/2025.03.12.642038

**Authors:** Natalie J. Tatum, Rebecca Scott, Gina M. Doody, Ian Hickson, Claire E. Jennings, Mathew P. Martin, Reuben M. Tooze, Julie A. Tucker, Anita Wittner, Lan-Zhen Wang, Eleanor K. Wright, Stephen R. Wedge, Martin E.M. Noble

## Abstract

IRF4, a transcription factor in the interferon regulatory factor family, is a key regulator in immune cell differentiation indicated to have an essential role in the development of lymphoid malignancies. Genome-wide association studies previously identified a set of overlapping mutations within the IRF4 DNA-binding domain in T-cell lymphoma and multiple myeloma, several of which appeared to be associated with better prognosis. Mapping these mutations to the known crystal structure of the IRF4:PU.1:DNA ternary complex and a new structure of the IRF4 DNA-binding domain in the apo state suggested they might interfere with DNA-binding, directly or via destabilisation of domain structure. We characterised these cancer-associated IRF4 mutants experimentally using the recombinant IRF4 DNA-binding domain (DBD) *in vitro* and examined the clinically relevant mutant K123R *in cellulo*. Using fluorescence polarisation, surface plasmon resonance, differential scanning fluorimetry and molecular dynamics, we find that mutation may give rise to significant differences in DNA-binding kinetics and thermal stability without compromising the affinity of IRF4 DNA-binding. The K123R IRF4 mutant showed increased transcriptional activity via a luciferase reporter assay and increased nuclear partitioning, which may be preferentially selected for in multiple myeloma. We discuss our observations in relation to the improved prognosis conferred by this mutation.

## Introduction

Interferon regulatory factor 4 (IRF4) is a transcription factor critical to the immune system, controlling a gene expression profile for differentiation of various lymphoid(1) and myeloid cell types – most notably the differentiation of mature B cells into immunoglobulin-secreting plasma cells.(2) As a direct result of this role, IRF4 has been associated with a variety of immune cell malignancies and evidence suggests certain subtypes of T cell lymphoma (TCL)(3) and multiple myeloma (MM)(4) are addicted to IRF4 expression programs, presenting a potential “Achilles’ heel” for exploitation in therapy. The role and function of IRF4 was reviewed by Agnarelli and colleagues.(5) IRF4 directly increases transcription of *MYC*, (4, 6) with significant downregulation of MYC target genes being observed upon IRF4 gene-silencing in anaplastic large cell lymphoma, (3) regardless of known driver anaplastic lymphoma kinase (ALK) fusion subtype. Moreover, microRNA-mediated knockdown of IRF4 has been shown in mouse xenograft models of MM to suppress tumour growth and proliferation, (7) further suggesting IRF4 silencing may be of therapeutic benefit. A spectrum of IRF4 mutations have been identified via genome-wide association studies (GWAS) within MM and TCL, (8–10) as well as loss/mutations in DLBCL(11) and IRF4 translocations within large B-cell lymphomas. Single-nucleotide polymorphisms within IRF4 have also been identified as conferring potential susceptibility to chronic lymphocytic leukaemia (CLL), (12) where mutations have been observed to enhance DNA binding, (13) and a recent study showed the C99R mutant, observed particularly in classic Hodgkin lymphoma, to have altered DNA-binding and induce a significantly modified gene expression programme.(14)

Particularly in MM, such IRF4 mutations are proposed to be activating or gain-of-function, in newly diagnosed MM conferring an improved outcome to triplet immunomodulatory drug treatment.(8) However, these mutations were also reported to be associated with a reduction in MYC expression, (8) and our mapping of the mutations onto the ternary complex structure of the murine IRF4 DNA-binding domain (DBD) with the PU.1 DBD and DNA(15) suggested that the majority could interfere with DNA binding or destabilise the domain’s structure. Given the uncertainties surrounding the functional consequences of these IRF4 mutations further characterisation is warranted.

IRF4 has two functional domains: a C-terminal interferon association domain (IAD) which mediates protein-protein interactions, and an N-terminal DNA-binding domain (DBD, **figure 1a**). IRF4, and the closely related IRF8, are unique within the IRF family in that it is not regulated by interferons via the IAD, rather it possesses an auto-inhibitory region within the C-terminus. This auto-inhibitory function is believed to regulate DNA interactions, physically preventing DNA-binding by IRF4 unless bound to a partner protein (such as PU.1), though the exact mechanism has yet to be elucidated.(16–18) However, a truncated mutant of IRF4 lacking the terminal 30 amino acids of the IAD has been demonstrated as hyperactive, leading to increased Th17 differentiation.(19) Members of the IRF family of transcription factors bind interferon-sensitive response elements (IREs), present in the regulatory regions of target genes, and are recruited and activated by a variety of co-factors to control transcriptional programs. Partners for IRF4 include PU.1, (20) Ikaros, (21), SPIB(22) and BATF, (23) but IRF4 can also act as a homodimer and notably PU.1 is down-regulated in myeloma cells.(5, 24)

**Figure 1.**
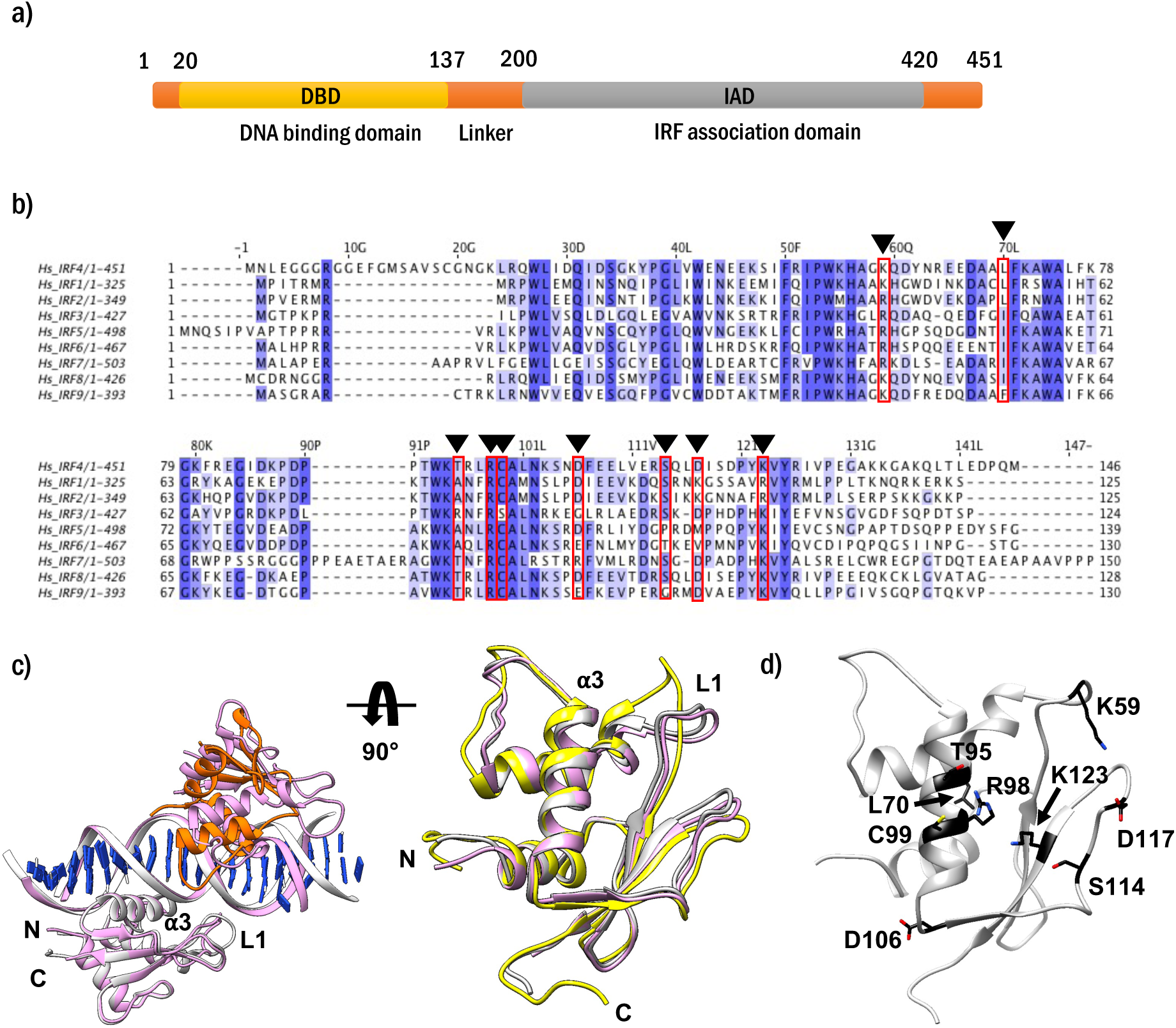
**(a)** Schematic of IRF4 domain structure with residue number boundaries indicated. **(b)** Comparison of sequences across the Interferon Regulatory Factor family. Sequence alignment created in JalView using ClustalW Multiple Sequence Alignment to *Homo sapiens* IRF4 sequence (Q15306). IRF sequences derived from UniProt. Residues coloured by conservation (%). Showing DBD and inter-domain ‘hinge’ region to residue 214 of IRF4. Mutated residues of interest are denoted in the wild-type sequence of IRF4. **(c)** Left panel shows the superposition, based on IRF4, of the IRF4:PU.1:DNA crystal structure (IRF4 and DNA in grey with PU.1 in orange) with the IRF4:DNA homodimer structure (PDB: 7JM4, shown in pink); right panel shows the superposition of the PU.1-bound IRF4 (grey), homodimeric IRF4 (pink, chain A) and first model of the murine apo NMR solution structure of IRF4 (PDB: 2DLL, yellow), with the DNA-binding ⍺3 helix and flexible L1 loop annotated. **(d)** Mutated residues (in black, labelled) mapped onto the human wild-type structure of IRF4 from the PU.1-bound structure. Figures prepared using Chimera.

The DBD is homologous across the IRF family, differing from a traditional winged-helix DNA-binding domain architecture in IRF4 by the insertion of two large internal loops capable of mediating further DNA interactions at the backbone, as observed in the crystal structure of the murine IRF4 DBD with the PU.1 DBD and a composite Ets-IRE DNA recognition sequence, and the NMR solution structure of the murine IRF4 DBD alone(15, 25). The core fold is conserved among the structures of other IRF family DBDs which have been elucidated in the apo form: the human IRF3 and mouse IRF7 structures were solved by X-ray crystallography;(26) the structure of mouse IRF2 was determined by NMR.(27) IRF7 is the most divergent, with a longer DNA-binding helix (**figure 1b**). A crystal structure of the human IRF4 DBD bound to both major grooves of an overlapping consensus ISRE DNA sequence in a homodimeric fashion has been reported.(13) Comparisons of apo and complexed forms of IRF4 showed that the largest differences corresponded to flexible loop L1 (residues 57-64) with minimal structural rearrangement of the core of the DBD (**figure 1c** and (13)).

Residues R98 and C99 within the DNA-binding helix of IRF4 (**figure 1b**) appear to be required for its recognition of a 5’-GAAA-3’motif in IREs(10, 15). The R98 residue is responsible for a base-specific interaction with DNA and is conserved across the IRF family. Many residues in this region of the protein are also conserved in identity or similarity to support protein-DNA contacts, for example, residue 99 is conserved as a cysteine in all but IRF3, where it is replaced by a serine. In IRF4, several of these residues are noted as mutational hotspots: K123 was present in 1.7% of cases of an MM cohort(8), mutations at K59, T95 and D106 have been found in TCL and MM, and K59R was observed in 6.1% of an adult TCL cohort.(10) In an adult TCL cohort an L70V mutation was observed in 3.3% cases, and mutations to residue S114 were also identified. S114 is on the PU.1 interacting loop as seen in the crystal structure of the ternary complex.(15) Previous studies have identified mutations to this loop, such as D117, which reduced ternary complex formation by up to 90%(30) and prevented rescue of IRF4 knockdown in diffuse large B-cell lymphoma cells, indicating mutations to the interacting loop region disfavour heterodimerisation.

Comparing the IRF4 sequence to other members of the family (**figure 1b**) suggests that some of these MM hotspot mutations should not impact DNA binding by IRF4. As previously stated, R98 and C99 are conserved while others diverge considerably from conserved residue identities. K123 is almost invariably a lysine, except where it is an arginine in IRF1 and IRF2; K59 is consistently a basic residue. L70 is typically a leucine, isoleucine, or valine across the family; mutation to a smaller hydrophobic residue may disrupt the hydrophobic core of the protein (**figure 1d**). T95 in IRF4 is either threonine or alanine in other IRFs, except in IRF3, where it is an arginine. The D106 residue is a glutamate in IRF6 and IRF9, an arginine in IRF7 and a glycine in IRF3, suggesting a charged residue is preferred but not essential at this position. The S114 and D117 residues are most divergent across the family, as would be expected for a flexible loop understood to mediate co-factor interactions. Moreover, some of these cancer-associated mutations may have roles in nuclear localisation, (31) ubiquitination(32) and phosphorylation, (33) when compared to equivalent positions in other IRF family members.

Herein we report the conformation of the human apo IRF4 through crystallisation of a surface entropy mutant, showing flexibility in the L1 loop and a stable core congruent with observations from the NMR and crystallised IRF4 complex structures.

To investigate the effect of the cancer-associated DBD mutations and characterise the IRF4-DNA interactions, we recombinantly expressed in bacteria the wild-type and a suite of mutant DBDs. Using fluorescence polarisation (FP), thermal melt, and surface plasmon resonance (SPR) assays, paired with molecular dynamics simulations, we then profiled their integrity and binding to a hairpin duplex IRE DNA consisting of only one canonical IRF4-binding site.

Further, using recombinant human protein expression systems we investigated the potential functional differences of select mutants of IRF4, and characterised cellular localisation and functional output using a reporter assay system. Our results suggest that the K123R mutation confers increased transcriptional activity with a skew toward increased nuclear residence indicative of an activating mutation, which may be preferentially selected for in MM.

## Materials and Methods

### General

Oligonucleotides for cloning were purchased from Eurofins Genomics and were solubilised in DNAse/RNAse-free water according to the manufacturer’s instructions. DNA sequencing was provided by Eurofins Genomics using standard primers (T7term for pBDDP vector; pShuttleCMV-F for pIRES2.EGFP.IRF4 vector). The pRL Renilla Luciferase vector (Promega) was used as a control for reporter assays. Purified plasmids were prepared from overnight cultures of *Escherichia coli* DH5α using QIAGEN plasmid spin miniprep kits according to the manufacturer’s instructions.

For protein purification, Ni^2+^-NTA resin and HiTrap FF columns were purchased from GE Healthcare and used according to the manufacturer’s instructions. 12% polyacrylamide Mini-PROTEAN® precast gels, and TGS running buffer (BIO-RAD), InstantBlue Coomassie stain (Expedeon) and PageRuler™ pre-stained protein ladder (ThermoFisher Scientific) were used for SDS-PAGE. General chemical stocks were purchased from Sigma-Aldrich.

### Mutagenesis, expression, and purification of mutants for assay

Fifteen mutants were generated by PCR mutagenesis using overlapping primers (**supplementary table S6**) and the plasmid, pIRES2.EGFP.IRF4 (a gift from Reuben Tooze and Gina Doody, University of Leeds), (22) which encodes human IRF4 (isoform 1) with an N-terminal Myc-epitope tag, as the template. DNA encoding the DNA-binding domain (residues 20-132) for sequence-validated mutants was amplified using the following oligonucleotides; forward primer (IRF4_G20_F2) 5’-TTTTCAGGGCGGATCCGGCAACGGGAAGCTCCGCCAG-3’ and reverse primer (IRF4_R11), 5’-GGTGGTGGTGCTCGACTAGGCTCCCTCAGGAACAATCCTGTAC-3’. The Cloning Enhancer (Clontech)-treated PCR product was then inserted into BamHI/XhoI-digested pBDDP-MBP by InFusion cloning (Clontech). The expression vector, pBDDP-MBP, (34) was a gift from Chris Gray (Beatson Institute, Glasgow). Digestion with BamHI and XhoI removes the sequence encoding the MBP tag and appends a His8-linker-His8-TEV sequence (**supplementary note S1**) to the N-terminus of the inserted PCR fragment. The K123R mutant was generated by PCR mutagenesis using the expression vector (pBDDP) containing the wild-type IRF4 DNA-binding domain (residues 20-132) as template and overlapping primers (**supplementary table S6**).

Sequence-validated expression vectors encoding IRF4 (20–132) with an N-terminal TEV-cleavable His8-His8-tag were transformed into *E. coli* Rosetta (DE3) pLysS. A 5 mL overnight starter culture in Luria Broth (LB) (Invitrogen) grown from a single colony, supplemented with kanamycin and chloramphenicol was used to inoculate 1 L of autoinduction media (Formedium) supplemented with 50 µg/mL kanamycin and 35 µg/mL chloramphenicol. The cultures were grown at 30°C for 24 hours in a shaking incubator (180 rpm). Cells were harvested by centrifugation (4000 x *g*, 4°C, 20 mins) and the pellet resuspended in 30 mL lysis buffer (25 mM HEPES, pH 7.0, 100 mM NaCl, 0.5 mM TCEP). Resuspended cells were frozen and stored at -20°C.

Cells were lysed by sonication on ice (10 minutes, 20 s on/40 s off, 30% amplitude) and the lysate clarified by centrifugation (1 hour, 48384 x *g*, 4°C). The supernatant was recovered, and His-tagged IRF4 purified by immobilised metal affinity chromatography (IMAC) using a 5 mL HisTrap column (GE Healthcare) and an ÄKTA Pure chromatography system (GE Healthcare) at 4°C. The column was washed with buffer A (40 mM HEPES pH 7.0, 300 mM NaCl, 0.5 mM TCEP, 20 mM imidazole) and eluted with a linear gradient to 100 % buffer B (40 mM HEPES pH 7.0, 300 mM NaCl, 0.5 mM TCEP, 500 mM imidazole). Fractions containing IRF4 were pooled and treated with recombinant His-tagged TEV protease at a ratio of 1:10 (protease:IRF4), overnight at 4°C. Each protein was then concentrated to 5 mL (Amicon Ultra 15 centrifugal filters, Ultracel 3K) and further purified by size-exclusion chromatography using a Superdex 75 16/600 column (GE Healthcare) equilibrated and run in phosphate-buffered saline (PBS) supplemented with 0.5 mM TCEP. A final subtractive IMAC polishing step was carried out using 4 mL NiNTA in a gravity flow column at 4°C. The column was equilibrated and run in PBS supplemented with 0.5 mM TCEP and 20 mM imidazole. 20 mM imidazole (pH 7.0) was also added to the load to ensure flow through of untagged IRF4. Fractions were checked for purity by SDS-PAGE. The purified IRF4 proteins were snap frozen in liquid nitrogen and stored at -80 °C. A sample of each batch of each mutant was subjected to LC-MS analysis to confirm identity and purity, with yields ranging from 1 – 5 mg/L. Protein concentrations were determined using a Thermo Scientific™ NanoDrop™ spectrophotometer, referenced against an appropriate buffer.

### Mutagenesis and sub-cloning of mutants for crystallisation and mammalian expression

The C99S and the E45A/E46A/K47A mutants of IRF4 were generated by PCR mutagenesis using overlapping primers (**supplementary table S6**) and the plasmid, pIRES2.EGFP.IRF4 as template. The E45A/E46A/K47A/C99S mutant was generated by sequential PCR mutagenesis using overlapping primers encoding the triple mutation (E45A/E46A/K47A) and pIRES2.EGFP.IRF4 as template, and then the overlapping primers encoding the C99S mutation and pIRES2.EGFP.IRF4 [E45A/E46A/K47A] as template. The DNA encoding the DNA-binding domain (residues 20-132) for sequence-validated mutants was then amplified using the following oligonucleotides; forward primer (IRF4_G20_F1) 5’-TTCCAGGGGC<u>CCATGG</u>GC<u>CAT*ATG*</u>GGCAACGGGAAGCTCCGCCAG-3’ and reverse primer (IRF4_R11), 5’-GGTGGTGGTGCTCGACTAGGCTCCCTCAGGAACAATCCTGTAC-3’. PCR products were subjected to PCR purification (QIAGEN PCR purification kit) before insertion into NcoI/SpeI-digested pET3dM-GST-3C-CCNT1(1–c259) by InFusion cloning (Clontech). The expression vector, pET3dM-GST-3C-CCNT1(1–c259), a derivative of pET3d (Novagen), was a gift from Richard Heath (Translational and Clinical Research Institute, Newcastle). Digestion with NcoI and SpeI removes the sequence encoding the CCNT1 insert and appends a GST-3C sequence (**supplementary note S1**) to the N-terminus of the inserted PCR fragment.

### Expression and purification of IRF4 surface-entropy mutant for crystallisation (IRF4 (20-132 [E45A E46A K47A C99S])

Plasmid encoding IRF4 (20-132 [E45A E46A K47A C99S]) with an N-terminal 3C-protease cleavable GST-tag was transformed into *E. coli* Rosetta (DE3) pLysS for expression. Several colonies were picked into a 60 mL LB over-night starter culture, which was used to inoculate 1 L of autoinduction media supplemented with 50 µg/mL ampicillin. The culture was grown at 37°C for 4 hours in a shaking incubator, then the temperature reduced to 18°C and growth continued overnight. The cells were harvested by centrifugation (4000 x *g*, 4°C, 20 minutes), and the pellet resuspended in 30 mL mHBS (25 mM HEPES, pH 7.4, 100 mM NaCl, 1 mM DTT). Resuspended cell paste was frozen and stored at -20°C.

Cells were lysed by sonication on ice for a total of 2 x 5 minutes (20 seconds on, 40 seconds off, 30% amplitude) and the lysate clarified by centrifugation (1 hour, 48384 x *g*, 4°C). The supernatant was recovered and filtered (0.45 µm). The clarified lysate was mixed with 5 mL glutathione Sepharose 4B (GE Healthcare) and incubated for 1 hour to overnight at 4°C on a roller. The mixture was poured into a gravity flow column, and the resin washed four times with four to five column volumes each time of mHBS before elution of bound protein with 20 mM glutathione in mHBS. The GST-tag was cleaved using GST-tagged 3C protease at a ratio of 1:40 (protease:fusion) overnight at 4°C. The cleaved eluate was concentrated to 2 mL and further purified by size exclusion chromatography using a Superdex 75 16/600 column (GE Healthcare), pre-equilibrated and run in mHBS. Fractions were checked for purity by SDS-PAGE, and those containing IRF4 were pooled. A final polishing step with a subtractive GSTrap column (GE Healthcare) was carried out. IRF4 was collected in the flow-through. The pure IRF4 was concentrated to 17 mg/mL, snap-frozen in liquid nitrogen and stored at -80° for later use (final yield approximately 1.4 mg/L).

### LC-MS characterisation

Intact protein masses were verified using an Agilent 6530 Accurate Mass Q-TOF instrument coupled to an Agilent 1260 Infinity II LC system. Purified protein (∼1 mg/mL, 50 μL) was desalted using Zeba Spin Desalting Columns (Thermo Scientific, PN: 10056033, 0.5 mL, 7 kDa MWCO) according to manufacturer’s guidelines. 20 μL of desalted protein was injected onto a Zorbax 300Å Stable Bond C8 column (Agilent, PN: 865973-906, 4.6 x 50 mm, 3.5 μm) for reversed phase separation at 60°C and 0.4 mL/min. Mobile phase was 0.1% (v/v) formic acid in LC-MS grade water (A) and acetonitrile (B) with separation performed over a linear gradient of 15-100% B over 25 min, followed by equilibration at 15% B for 10 min. Proteins were detected in positive ion mode using electrospray ionisation with nebuliser pressure of 45 psi, drying gas flow of 5 L/min and source gas temperature of 325°C. Sheath gas temperature of 400°C and gas flow of 11 L/min, capillary voltage of 1750V and nozzle voltage of 2000V were also applied. Mass spectra were acquired using Mass Hunter Acquisition software (version B.08.00) over 100-3000 m/z range, at a rate of 1 spectra/s and 1000 ms/spectrum, using extended dynamic range mode (2 GHz). Subsequent intact mass determination was achieved using Agilent Mass Hunter Bioconfirm software (version B.08.00) with maximum entropy deconvolution. ESI-QTOF raw data and deconvoluted mass spectra for WT, K123R (and K59R) IRF4, are given as **supplementary figure S9**; protein intact masses for all recombinantly expressed mutants and wild type IRF4 ± error are given in **supplementary table S2**.

### FP assay development, optimisation and data generation

Oligomeric DNA hairpins for assay were designed using OligoAnalyzer 3.1 from IDT (www.idtdna.com/calc/analyzer). Care was taken to ensure only one predicted structural outcome with the desired four-nucleotide hairpin turn and a predicted Tm > 65°C. The resulting designed oligomers were sourced from Eurofins Genomics with a 3’ fluorescein (FAM) label. Upon receipt, lyophilised hairpin DNA oligomers were re-suspended to a concentration of 100 µM in DNAse/RNAse-free water and stored at -20°C. For use, DNA was defrosted and a 5 µM solution prepared in an amber Eppendorf tube to prevent any degradation of the label. DNA was heated to 98°C for 5 to 10 mins, and the solution was allowed to cool to room temperature to anneal the hairpin. Annealed DNA was stored at 4°C for further use.

The fluorescence polarisation assay was optimised for 20 µL volume. Hairpin DNA concentration for assay was determined by serial dilution from 125 – 0.002 nM in PBS, with assay concentration set at maximum signal for minimal concentration, in this case 5 nM. Protein was titrated against a fixed concentration of DNA from a top concentration of 10 µM protein in an 11-point 3-fold serial dilution in PBS supplemented with 0.05% Tween20. FAM-labelled hairpin duplex at a final concentration of 5 nM was added to each well to give a final assay volume of 20 µL. The plate was incubated for 1 hour at room temperature before being read by a BMG Labtech PHERAstar FS microplate reader, also at room temperature. Each well was subject to 200 flashes of light at excitation wavelength 485 nm, with emission at 520 nm recorded. A positioning delay of 0.3 s was used and multiple rows were read in serpentine fashion. Gains and focal height were adjusted per plate based on a reference well of free fluorophore at fixed assay concentration (5 nM). Binding curves were analysed in GraphPad Prism 6 and fit to a one-state binding model with Hill slope; all curves are combined and presented as **supplementary figure S10** with error bars representing range.

### SPR assay development and data generation

All SPR binding experiments were performed on a Biacore® S200 (GE Healthcare, Uppsala, Sweden) instrument at 20 °C in a running buffer composed of PBS, 0.05 % (v/v) Tween20, 0.25 mM TCEP, pH 7.4, at a flow rate of 30 μL min^-1^, 30s contact and 90s dissociation time. Running buffer was prepared freshly on a weekly basis and filtered (0.22 μm, Whatman cellulose nitrate filters) and degassed prior to SPR analysis. The sample compartment was held at 4°C to prolong the lifetime of the protein samples.

3’-biotinylated oligonucleotide hairpins were captured on streptavidin pre-coated SA sensor chips (GE Healthcare, Uppsala, Sweden). The streptavidin (SA) sensor surface was first conditioned with three consecutive 1-min injections of high salt solution (50 mM NaOH in 1 M NaCl) followed by a wash of the same plus 50% isopropanol. Next, annealed biotinylated hairpin oligonucleotides were diluted in running buffer 1000-fold from a 0.1 mM stock solution, and then diluted 250-fold to a final concentration of ∼0.4 nM. Sequential pulses of biotinylated oligonucleotides were applied over the streptavidin sensor surface at a flow rate of 10 μL min^-1^ to achieve immobilization levels of 41-58 response units (RU). Injection lines were washed with isopropanol between injections. Oligonucleotides were immobilized freshly upon significant degradation of maximal response in the presence of IRF4 (wild-type) as analyte. Free biotin was not applied to the sensor surface after ligand capture as this is not recommended for the Biacore® S200.

Initial experiments used free streptavidin (flow cell 1) as the reference surface with IRE hairpin captured on flow cell 2. Latterly, a scrambled oligonucleotide (scrIRE) was designed and captured on flow cell 3 for comparison against free streptavidin on flow cell 1 and IRE on flow cells 2 and 4.

Purified IRF4 mutants were exchanged into freshly prepared SPR running buffer using PD10 desalting columns (GE Healthcare), and then concentrated using centrifugal concentrators (Amicon Ultra 0.5 mL, Ultracel 3K MWCO) to a concentration sufficient for each experiment. A 1 M NaCl wash step was applied between each injection to remove residual bound protein from the chip, 30 μL min^-1^ for 30 s.

All SPR data processing and analyses were performed using BiaEvaluation Software (version 1.0) and GraphPad Prism 6. All monitored binding resonance signals were double-referenced, *i*.*e*., signals monitored on the reference channel were subtracted from signals monitored on the liganded channels, which were then further adjusted to allow for buffer controls (**supplementary figure S11**). For kinetic evaluation, data were globally fit to the mathematical binding model describing either a one-state interaction (**supplementary table S5**) or a two-state interaction (**supplementary table S4**). For steady state affinity analyses, the SPR signals at equilibrium were plotted against analyte concentration and fit to the one-to-one steady state interaction model with four parameters. All three derived K_D_ values are presented herein.

### Molecular dynamics simulations and MM/PBSA calculations

The starting structure for molecular dynamics simulation (MD) was derived from the crystal structure of the ternary DNA-IRF4-PU.1 complex(15) (AK Aggarwal, co-ordinates supplied by personal communication). The PU.1 chain was removed along with all water and solvent molecules. The wild-type human DNA-IRF4 sequence was achieved by reversion of residue V49 to I49. This and subsequent mutants were derived *in silico* by altering the respective residues to the lowest energy rotamer in UCSF Chimera.(35)

Simulations were performed in GROMACS 5.1.4 using one NVIDIA GeForce GTX 1080 GPU and ten Intel Xenon E5-2630 CPU threads. Each system was parameterised in the CHARMM27 forcefield(36, 37) and solvated in a cubic box with a 10 Å shell of TIP3P water.(38) The systems were then neutralised using NaCl to a final concentration of 0.1 M, followed by steepest descent energy minimisation over 5000 steps. Each system was subject to 200 ps equilibration at 300 K in the NVT ensemble (2 fs steps with Nose-Hoover temperature coupling). Equilibration was continued in the NPT ensemble for 500 ps using 2 fs steps and Parrinello-Rahman pressure coupling. Position restraints were used throughout equilibration and released for the production MD, consisting of 8 ns simulation time in the NPT ensemble, conducted in triplicate from the equilibrated structure. Energies and co-ordinates were written every 5000 steps (10 ps) for analysis. Convergence was determined by tracking the Lennard-Jones and coulombic energies throughout the simulation, as well as the root-mean-square deviation (RMSD) of the protein alpha-carbons. All 54 simulations in total were suitable for use in MM/PBSA calculations.

GROMACS functions were used to analyse trajectory RMSDs and per-residue root-mean-square fluctuations (RMSFs). The final, converged 2 ns (6 – 8ns, total 201 snapshots at 10 ps intervals) of each simulation was used in a single-trajectory approach to calculate binding energies using the MM/PBSA method with the *g_mmpbsa* package.(39) A solute dielectric constant of 4 was used to reflect the presence of DNA in the complex. Output binding energies from the three independent wild-type simulations were averaged to provide a baseline for ΔΔG calculations. Binding energies were plotted against averaged experimental kinetic measurements derived from SPR experiments described above (**supplementary figure S12**).

### Differential scanning fluorimetry

Optimal conditions for differential scanning fluorimetry (DSF) were ascertained using wild type IRF4. Protein concentrations from 60 µM to 7.5 µM and SYPRO Orange concentrations from 10X to 2X were investigated to optimise signal-to-noise. A final concentration of 60 µM protein with 10X SYPRO Orange was used for all other DSF experiments. Proteins were freshly buffer exchanged into PBS supplemented with 0.25 mM TCEP for each experiment.

14 µL of 64 µM protein were aliquoted in triplicate into a 384-well plate, and 1 µL 150X SYPRO Orange was dispensed into wells to give a final concentration of 60 µM protein and 10X SYPRO Orange and a final volume of 15 µL. Plates were immediately analysed using a Via7 Real-Time PCR System (Applied Biosystems). Samples were equilibrated at 25°C for 2 mins, and then the temperature was increased at a rate of 0.05°C/s to a final temperature of 99°C, which was held for a further 2 mins. The ROX reporter was used with quencher and no passive reference; the melt curve optical filter selected was x1-m3. Data were analysed using the Protein Thermal Shift software v1.3 (Applied Biosystems). Measurements were made on two independently prepared batches of protein.

### Statistical methodology

Statistical significance of results for DSF, SPR, FP and MM/PBSA was determined in all cases by unpaired parametric t-test with Welch’s correction (n.s. = not significant, * p=0.0332, ** p=0.0021, *** p=0.0002, **** p=<0.0001). Statistics for luciferase reporter assay were calculated using two-tailed unpaired Student’s t-test of unequal variance using Welch’s correction, with p<0.05 of statistical significance.

### Crystallisation of IRF4 DBD in the apo state

Initial sparse matrix and grid screens (Hampton Index HT) were set up using a Mosquito robot (TTP Labtech) to set drops of 300 and 450 nL final volume at 1:1 and 2:1 protein:reservoir ratios, respectively, where protein was at 14 mg/mL in mHBS in SwissCI 2-well sitting drop plates, which were then sealed with optically clear tape and incubated at 277 K and 293 K. Crystals were observed at 277 K after one week in condition G8 (25 % PEG-3350, 0.1 M HEPES pH 7.5, 0.2 M ammonium acetate). Crystals were harvested in mother liquor supplemented with 20% glycerol and flash frozen in liquid nitrogen for diffraction testing and data collection on an in-house rotating anode source (Rigaku MicroMax007) equipped with an R-Axis IV image plate detector (Rigaku) and an Oxford CryoSystems cryo-stream. A partial dataset was collected at 100 K to 2.3 Å, and indexed in space group C222_1_ with cell dimensions 64.7, 66.6, 68.6 Å.

Grid screens at 277 K around these conditions sampling pH and % PEG3350 against fixed salt concentration (either 0.2 M NaCl or 0.2 M ammonium acetate), yielded multiple hits, however, all crystals were layered in appearance and showed evidence of OD twinning in the diffraction pattern.

Optimised crystals for seeding were grown at 277 K by sitting drop vapour diffusion by mixing protein in mHBS at 14 mg/mL with reservoir solution (0.1 M HEPES pH 7.5, 25 % PEG 3350, 0.2 M ammonium acetate) at a ratio of 2:1 to make a 450 nL drop. Crystals grew to final size within 3 days. Seeds were prepared by addition of 5 μL stabilising solution (0.1 M HEPES pH 7.5, 27.5 % PEG 3350, 0.2 M ammonium acetate, 0.1 M NaCl) to the drop, which was then taken up into a micro-centrifuge tube containing 20 μL stabilising solution and a Teflon bead. The cover slip was rinsed twice with 5 μL stabilising solution, and the total volume made up to 50 μL. The sample was vortexed for a total of 4 mins (in 30 secs bursts interspersed with 30 secs on ice). Seeds were then diluted to ∼18 % v/v into stabilising solution and flash frozen in liquid nitrogen before storage at 193 K.

Random micro-seed matrix screening into Hampton Index HT at 293 K gave single, diffracting crystals in multiple conditions. All crystals showing sufficient diffraction on a rotating anode source could be indexed in space group C222_1_. Conditions D8 (0.1 M HEPES pH 7.5, 25 % w/v PEG3350) and E6 (30 % PEG550MME, 0.1 M Bis-Tris pH 6.5, 0.05 M CaCl_2_) were selected for further work as they gave good quality crystals that could be harvested directly from the drop without the need for additional cryo-protectant.

Optimised crystals were grown in SwissCI 3-well sitting drop plates at 293 K. Drops were set using a Mosquito with a humidifying chamber set to high humidity, and a pre-chilled block for protein and seed dispensing. Wells were filled with 45 μL chilled reservoir solution (25 % PEG 3350, 0.1 M HEPES pH 7.5). Protein and seeds were defrosted from snap-frozen stocks on ice, and protein was spun for at least 10 mins (16000 x *g*, 277 K). Drops were dispensed as follows; 300 nL IRF4 (20-132 [E45A E46A K47A C99S]) at 14 mg/mL, 50 nL seed stock, 100 nL reservoir solution; the plate was sealed with optically clear film and stored at 293 K. Crystals began to appear almost immediately and reached final size within two days.

### Data collection and processing, structure solution and crystallographic refinement

Three hours prior to data collection, 50 nL DMSO was dispensed into the drop and the plate re-sealed. Crystals were harvested directly into liquid nitrogen without further cryo-protection. Data were collected at 100 K on beamline I04-1 at the Diamond Light Source. Data collection and refinement statistics are given in **supplementary table S3**.

Data were processed using the automated software pipeline available on the beamline at the Diamond Light Source. The structure was solved by molecular replacement using a partially refined structure of the human IRF4 DBD derived from an incomplete dataset at 2.3 Å, and initial rounds of refinement were carried out automatically using AutoSolve.(40) Manual model completion was carried out in Coot(41) interspersed with rounds of reciprocal space refinement in Buster(42), Refmac5(43) and phenix.(44) The final model comprises residues 21 to 58 and 65 to 132 of the human IRF4 sequence with one additional C-terminal residue (L133) derived from the sub-cloning process. Residues 59 to 64 could not be modelled due to lack of electron density.

The IRF4 DNA-binding domain surface entropy mutant apo structure has been deposited in the Protein Structure Databank under accession code 6TD4.

### Mammalian cell expression, nuclear localisation and IRF4-dependent reporter assay

Trypsin-harvested cells were counted using Countess II FL automated cell counter (Thermofisher), seeded at 1.8×10^-5^ cells/mL in 6-well plates unless otherwise specified, and incubated for 24 h at 37⁰C. 0.75 µg of pIRES2.EGFP.XBP1S, pIRES2.EGFP.IRF4 full length WT, K123R, K59R and R98A/C99A plasmid were transfected into cells using a 3:1 ratio of TransIT-LT1 Transfection Reagent (Mirus) to plasmid in Opti-MEM®I Reduced-Serum Medium (Gibco) and incubated at 37°C for 24 h. For dual-luciferase assays, cells were also co-transfected with 0.75 µg pFireflyL (pGL4.10 (Promega), containing a 2.5 kb region upstream of the *ZBTB32* transcription start site harboring an AICE-1 type IRF element with the sequence 5’-GGAAGATGAGTCAGAACGAAAGGGAAAATG-3’, and 5 ng of pRenillaL. Non-transfected cells containing no plasmids were used as a negative control for IRF4/GFP expression.

0, 6 or 24 h after transfection, trypsin-harvested cells were centrifuged at 500 x *g* for 5 minutes, washed in PBS and proteins extracted as whole cell lysates or fractionated samples. For whole cell lysates, cells were lysed in Phosphosafe (Novagen) with Protease Inhibitor Cocktail (Roche), centrifuged at 16000 x *g* and 4°C for 10 minutes and supernatant collected. For subcellular fractionation, NE-PER™ Nuclear and Cytoplasmic Extraction Reagents (Thermofisher) were used according to the manufacturer’s instructions, with an extra wash with Cytoplasmic Extraction Reagent I after cytoplasmic extraction.

Total protein concentration was measured by BCA assay (Pierce™ BCA, Thermofisher) against a BSA standard (0-1.2 mg/mL). 1-30 µg of protein was prepared with 4X Laemmli Buffer (Biorad) or XT buffer (Biorad), boiled at 95 °C for 5 minutes and loaded onto 4-20% Criterion TGX Precast Gel (Biorad) for electrophoresis in Tris-Glycine SDS running buffer at 100V for 2 h. Protein was transferred to nitrocellulose membrane (GE Lifesciences), 0.45 µm pore size, at 100V for 1 h in Tris-Glycine transfer buffer and stained with Ponceau-S (Sigma). Membranes were blocked with blocking buffer (5% (w/v) powdered skimmed milk in Tris-buffered saline supplemented with 0.05% Tween20 (TBST)) and incubated with primary antibodies in blocking buffer overnight at 4 ⁰C (Anti-MUM1, Ab133590, (Abcam); Anti-GFP, 66002-1-Ig (Protein-Tech); Anti-MYC, Sc-40, Anti-GAPDH, 03411 and Anti-PARP1, F-2 (all SantaCruz Biotechnology). Membranes were washed for 3 x 5 minutes in TBST, incubated with diluted secondary antibodies (Anti-mouse Ig-HRP, P0447; Anti-rabbit Ig-HRP, P0448 (Dako)) for 1 h and washed for 3 x 10 mins in TBST. Proteins were visualised with Clarity Western ECL Substrate (Biorad) using an Odyssey Fc imaging system (LI-COR) and quantified by Empiria Studio 1.1 (LI-COR).

HEK293T cells were co-transfected with 0.75 µg of pFireflyL, 5 ng pRenillaL, and 0.1 µg of pIRES2.EGFP.IRF4 WT, K123R or K59R plasmid. After 24 h, cells were scraped into passive lysis buffer and subjected to 2 freeze-thaw cycles for complete cell lysis. GFP fluorescence was evaluated by PHERAstar FS (BMG Labtech) and luciferase activity by Dual-luciferase Reporter Assay System (Promega) using FLUOstar Omega (BMG Labtech). Firefly luciferase activity was expressed as firefly/renilla activity, normalised against GFP expression. Values are expressed as mean ± standard error for n=5 measurements.

### Immunofluorescence

Methanol-cleaned dried coverslips were placed in wells before seeding with HEK293T, or HEK293T cells transfected with pIRES2.EGFP.XBPS1, WT or K123R IRF4. After 24 h, coverslips were washed in PBS and fixed in ice-cold 100% methanol for 1 h at -20 °C. Cells were permeabilised in 0.1% (v/v) PBS-TritonX (Sigma Life Science) and blocked for 1 h with 4% (v/v) goat serum (Dako) in 0.1% PBS-TritonX. Coverslips were incubated in primary antibodies overnight and washed 3 times in PBS before 2h dark incubation with secondary antibodies (Anti-rabbit Alexa Fluor 647, A21244; Anti-rabbit Alexa Fluor 594, A11037; Anti-mouse Alexa Fluor 488, A11029 (Invitrogen)). Coverslips were washed twice with PBS and once with ddH_2_O, dried then mounted on glass slides using ProLong™ Glass Antifade Mountant with NucBlue™ Stain (Invitrogen). Slides were visualised using a Leica TCS SPE confocal microscope and analysed with Fiji (ImageJ2, Madison, Wisconson, USA).

## Results

### Cancer-associated mutations in the IRF4 DBD map to key regions of the structure

Cancer-associated mutants K59N, K59R, L70V, T95A, T95K, T95R, C99R, D106Y, S114N, S114R and K123R were selected as known hotspot mutations in MM and TCL.(8–10)

Initially, we used the mutation prediction software packages MutationTaster, (45) PROVEAN, (46) SIFT(47) and CADD(48) to gauge the potential functional effects of these mutations (**supplementary table S1**). Interestingly, MutationTaster predicted all variants – including the natural variation between human and mouse IRF4 (I49 in human IRF4 is valine in murine IRF4) – to be ‘disease causing’ with high confidence. In contrast, PROVEAN suggested I49V, L70V and S114N to be neutral and the remainder to be deleterious; SIFT predicted a mix of tolerated and not tolerated variants; and CADD indicated most variants were in the top 1% most deleterious mutations. Overall, the analysis indicated most, if not all, variants would have some negative effect on IRF4 function.

We mapped the mutations to the IRF4:PU.1:DNA crystal structure and observed that many would be expected to disrupt DNA interactions or protein integrity. For example, K59, T95, D106, S114 and K123 lie on the surface of IRF4 close to the binding interface with DNA (**figure 1c-d**) and each of these residues lies within hydrogen bonding networks that could be disruptive to protein-DNA interactions if perturbed by mutation (**supplementary figures S1-5**). When mapped to the murine IRF4 apo solution NMR structure, (25) all but K59 reside in well-ordered parts of the ensemble; K59 sits on the highly flexible L1 loop. In contrast, L70 lies within the IRF4 DBD hydrophobic core and mutation would be expected to reduce stability of the fold through loss of contacts with W27 and I52 (**supplementary figure S6**). Finally, C99 lies on the IRF4:DNA interface and interacts with the phosphate backbone; depending upon the nature of the mutation, change at this position could be tolerated (*i.e.* C99S, which was generated for crystallisation), or disruptive (*i.e*. C99A or C99R**, supplementary figure S7**). The C99A mutant in particular has been reported to have reduced binding affinity for DNA.(17) Several engineered mutants from the literature were also selected for study: the R98A and the double mutant R98A/C99A were selected as the R98A mutant abolishes the base-specific interaction, and the R98A/C99A mutant has been demonstrated to abolish DNA binding;(17) the mutants D117A and D117H were reported to rescue IRF4 knockdown by shRNA.(30) Together, these engineered mutants represent both negative (R98A/C99A) and positive (D117A, D117H) control examples.

### The crystal structure of IRF4 in the apo state reveals conformational change in the DNA-binding site

To better understand the influence of DNA and co-factor binding on the structure of the IRF4 DBD, we undertook to determine the crystal structure in the absence of both DNA and PU.1. We designed a minimal construct comprising residues G20 to A132 based on the modelled residues in the IRF4-PU.1-DNA ternary complex structure, however, this protein failed to yield diffraction-quality crystals. The purified IRF4 DBD tended to form disulphide-linked dimers through its single cysteine residue, C99. To eliminate this source of heterogeneity we made a C99S mutant which showed improved solution behaviour, but also failed to yield diffraction-quality crystals. In order to improve crystallisation propensity, we used the SERp tool(49) to identify different combinations of surface mutations that might reduce the “surface entropy”(50) without impacting on the DNA-binding interface. Alanine mutation of a patch comprising residues E45, E46 and K47 resulted in a protein that crystallised more readily, however, these crystals were poorly ordered. Combination of C99S and E45A/E46A/K47A mutations yielded protein that gave crystals from an initial screen which diffracted to 2.3 Å on a rotating anode source. However, this crystal form proved challenging to reproduce, and subsequent crystals showed a layered structure under the microscope which translated into “OD twinning” in the diffraction pattern. In order to overcome these limitations, we utilised a random micro-seed matrix screening (rMMS) approach(51) to identify multiple novel crystallisation conditions, including two conditions that allowed crystals to be frozen directly without additional cryo-protectant. Crystals from both conditions gave diffraction to better than 2 Å on beamline I04-1 at the Diamond Light Source. We here describe the IRF4 DBD structure derived from a dataset to 1.7 Å collected from a seeded crystal grown from 0.2 M HEPES pH 7.5, 25 % PEG 3350.

The crystal structure of the human IRF4 DBD in the absence of DNA closely resembles the NMR ensemble (PDB 2DLL), and the crystal structure of murine IRF4 DBD in complex with the PU.1 DBD and DNA, (15) and comprises a four-stranded β-sheet flanked by three α-helices (a so-called ‘winged-helix’ architecture) (**figure 2a**). The major difference between the DNA-bound and apo structures lies in the DNA-binding loop (residues 54 to 67), which is highly mobile in the NMR ensemble, and could only be partially modelled in the crystal structure; residues 59 to 64 at the tip of the loop are not visible in the electron density. Comparison to the DNA-bound structures suggests this loop may undergo a transition to a more ordered state upon DNA-binding (*viz.* **figure 1c**; **figure 2d**).

**Figure 2.**
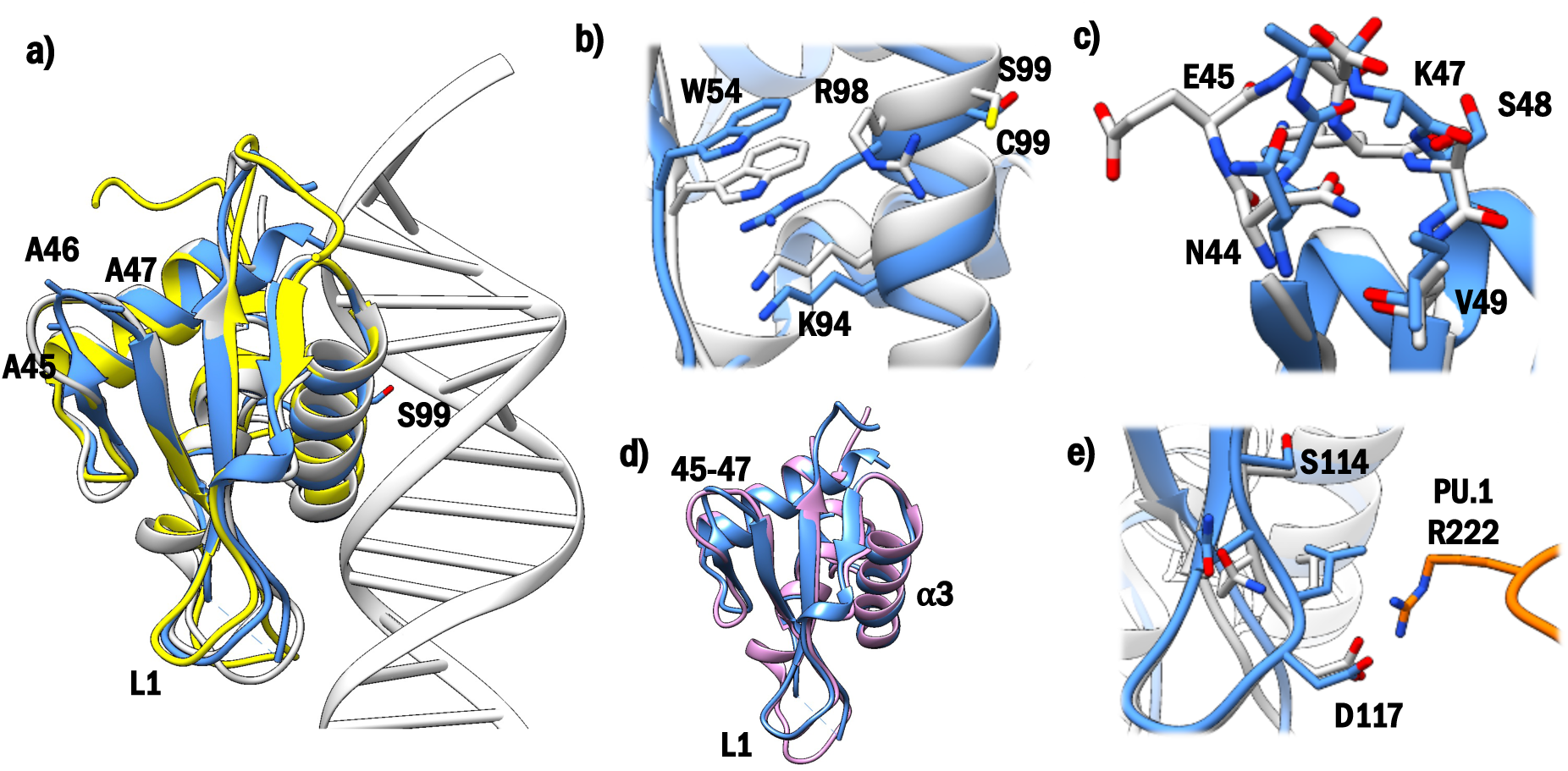
**(a)** The crystal structure of the human apo IRF4 DNA-binding domain (blue) closely resembles that of murine IRF4 in complex with DNA (both grey) and PU.1 (not shown) and the NMR structure of murine IRF4 (yellow, PDB: 2DLL). Side-chains of the surface residues mutated in the crystallised human IRF4 are shown as ball and stick and labelled. **(b)** Comparison of the region surrounding S99 in the human apo IRF4 DNA-binding domain (blue) with that of C99 in the murine IRF4 (grey) complex with DNA. In the absence of DNA R98 folds into the protein and intercalates between W54 and K94. The bound DNA is omitted for clarity. **(c)** The loop carrying the surface entropy reduction mutations adopts a different conformation in the human (blue) and murine (grey) IRF4 structures. **(d)** Superposition of the human apo IRF4 (blue) with the homodimerized IRF4 (PDB: 7JM4, pink), highlighting the more ordered L1 loop on DNA-binding and the alternate conformation at residues 45-47. **(e)** Comparison of the region surrounding D117 in the human apo IRF4 DNA-binding domain (blue) and the murine IRF4 (grey) complex with PU.1 (orange) and DNA. In the absence of DNA and PU.1, local side-chain conformations are preserved, however, the peptide backbone of the PU.1 binding loop moves away from its position in the complex. Structures were superimposed using the DNA-binding helix (residues 91-103). DNA omitted for clarity. Figures prepared using Chimera.

The engineered C99S mutation lies within the DNA-binding interface (**figure 2a**). A number of side-chain movements are observed in the vicinity of the engineered C99S mutation, including most notably R98, which rotates and extends to stack against the side-chain of W54 in the absence of DNA (**figure 2b**). A similar conformation of R98 is also seen in one of the 20 models within the NMR ensemble (PDB 2DLL model 16), where the side-chain of R98 within the ensemble shows considerable flexibility. The equivalent residue (R96) in IRF7 adopts an identical conformation in the apo crystal structure of the IRF7 DBD (PDB 3QU3).(26) In IRF2 (PDB 1IRF)(27) and IRF3 (PDB 3QU6), (26) the equivalent arginines also adopt different conformations in the free compared to the DNA-bound structures, although these differ from those observed in IRF4 and IRF7, and packing against tryptophan is not observed. This suggests that R98 conformational change upon DNA-binding may be a common feature across the IRF family, and in our IRF4 structure is not related to the C99S mutation *per se*.

The engineered surface-entropy-reduction mutations (E45A/E46A/K47A) lie on a surface-exposed loop connecting strands β1 and β2, which is not involved in contacts with either DNA or the PU.1 DBD (**figure 2a**). The backbone conformation of this loop differs from that observed in the wild-type structures and is stabilised by contacts between the side-chain CO of N44 and the backbone NH of A46 and A47 (**figure 2c**). In the wild-type murine IRF4 structure, N44 makes similar contacts to the backbone NH of K47 and S48. The single sequence difference between the human and murine IRF4 DBD, I49V, lies within this loop. The N44 side-chain as modelled in the murine crystal structure would generate a close contact with the additional methyl of I49 in the human crystal structure and is observed to do so in the NMR ensemble model (I49 Cδ1 to N29 Nδ2 distance of closest approach ranges from 2.9 to 3.4 Å). Thus the altered conformation of this loop likely arises due to the engineered (E45A/E46A/K47A) mutations, a conclusion supported by comparison to the homodimeric IRF4:DNA crystal structure, where the mutated loop deviates notably in conformation despite otherwise high structural similarity (RMSD 0.79 Å over 94 pruned Cα pairs, 1.58 Å over all 103 Cα pairs).

The region surrounding D117 comprising the interface with the PU.1 DBD shows limited change in local conformation, however, an increase in B-factor for this region suggests increased flexibility in the absence of DNA and PU.1. Overlay of the free and bound structures shows a shift of ∼ 1.5-1.7 Å in the backbone away from the DNA/PU.1-bound position (**figure 2e**). This shift is possibly a consequence of crystal contacts from a neighbouring molecule in the region surrounding S119 (data not shown).

Inspection of the regions surrounding the cancer-associated mutations characterised in this study shows that for L70, T95, D106, S114 and K123 the protein environment remains largely unchanged (data not shown), whilst K59 cannot be modelled due to the increased flexibility in the DNA-binding L1 loop in the apo state.

### Biophysical characterisation of IRF4 wild-type and mutant DBDs

To investigate the interactions of the IRF4 DBD with DNA, we utilised the minimal construct previously designed for crystallisation, having verified its integrity and structure by X-ray crystallography. We hypothesised that a subset of mutations in the IRF4 DBD may affect the packing of the hydrophobic core or otherwise induce conformational changes resulting in reduced protein thermostability (**figure 1c, supplementary figures S1-S7**). We therefore used differential scanning fluorimetry (DSF) to characterise the effect of mutations on protein melting temperature (Tm). The wild-type IRF4 DBD was determined to have a Tm of 58.6 ± 0.8 °C (± SD, **table 1**). Mean Tm for each of the mutant IRF4 proteins is presented in **table 1**, whilst **supplementary figure S8a** shows the experimental variation. Minor variations in protein melting temperature were measured across the mutant cohort, however, despite statistical significance in several cases, the observed melting temperatures are not likely to have physiological consequences, *i.e.* wild type and mutant melting temperatures are well in excess of physiological temperatures.

**Table 1:** Summary results for wild-type IRF4 and all mutants studied. Melting temperatures determined for IRF4 wild-type and cancer-associated mutants. Temperatures are given in degrees Celsius ± standard deviation from the mean. Binding affinities of IRF4 proteins were obtained using an FP assay and fitting to a one-site binding model (with Hill slope) in GraphPad Prism 6. IRF4 wild-type and mutant affinities for DNA as determined by SPR were fit using the steady state affinity model and a two-state kinetic binding model. Standard error (SD/√N) is given in parentheses for FP and steady-state SPR measurements. Measurements were made over two independently grown, expressed and purified protein batches.

| Protein | T<sub>m</sub> / °C | K<sub>d</sub> by FP/ nM | K<sub>D</sub> by SPR – Steady State Affinity / nM | K<sub>D</sub> by SPR – Kinetic Two-State Fit / nM |
| --- | --- | --- | --- | --- |
| Wild-type | 58.6 ± 0.8 (0.22) | 565 ± 191 (61) | 402 ± 165 (55) | 187 ± 143 |
| K59N | 57.4 ± 1.3 (0.30) | 6002 ± 3156 (1402) | > 4500 | 107 ± 133 |
| K59R | 58.5 ± 2.3 (0.55) | 350 ± 275 (112) | 275 ± 93 (33) | 460 ± 429 |
| L70V | 53.7 ± 3.3 (0.87) | 127 ± 27 (10) | 167 ± 82 (29) | 121 ± 111 |
| T95A | 58.9 ± 1.0 (0.27) | 105 ± 59 (34) | 186 ± 82 (31) | 305 ± 174 |
| T95K | 57.5 ± 0.5 (0.13) | 298 ± 154 (66) | 594 ± 251 (112) | 913 ± 147 |
| T95R | 59.3 ± 1.1 (0.27) | 151 ± 72 (41) | 449 ± 49 (25) | 481 ± 149 |
| R98A | 58.8 ± 1.9 (0.53) | > 10000 | > 4500 | > 4500 |
| R98A/C99A | 60.3 ± 1.2 (0.35) | > 10000 | > 4500 | > 4500 |
| C99A | 59.1 ± 1.5 (0.43) | 353 ± 169 (64) | 173 ± 60 (23) | 61 ± 29 |
| C99R | 56.8 ± 0.8 (0.22) | 279 ± 286 (143) | 546 ± 174 (87) | 660 ± 317 |
| C99S | 59.1 ± 0.5 (0.16) | 243 ± 238 (97) | 338 ± 173 (65) | 139 ± 43 |
| D106Y | 56.7 ± 3.2 (0.92) | 219 ± 261 (151) | 178 ± 43 (15) | 135 ± 121 |
| S114N | 54.9 ± 5.3 (1.52) | 181 ± 14 (7) | 291 ± 79 (28) | 145 ± 151 |
| S114R | 54.8 ± 4.0 (1.34) | 332 ± 162 (93) | 213 ± 94 (30) | 49 ± 26 |
| D117A | 56.7 ± 2.3 (0.68) | 109 ± 31 (18) | 301 ± 87 (33) | 293 ± 213 |
| D117H | 58.3 ± 0.9 (0.26) | 131 ± 28 (16) | 247 ± 84 (34) | 204 ± 131 |
| K123R | 58.2 ± 0.9 (0.22) | 143 ± 34 (17) | 491 ± 48 (20) | 1107 ± 341 |

Having thus established the thermostability of the IRF4 DBD mutants, we set out to functionally characterise cancer-associated IRF4 mutations by determining their effect on DNA-binding affinity. Fluorescence polarisation (FP) assays were utilised to measure the DNA-binding capability and affinities of each of the mutants, using a short DNA hairpin (IRE sequence: 5’-CTTGGTTTCACTTCCTGCGTTTTCGCAGGAAGTGAAACCAAG-3’) designed to fold back to form a double strand via the TTTT motif and based on the consensus sequence used in the ternary complex crystal structure of IRF4.PU-1.DNA(15), possessing the 5’-GAAA-3’ canonical IRF binding motif.

Proteins were titrated against the IRE DNA over a concentration range of 0.1 – 10, 000 nM; one-site binding models were fit to the dose-response curves (**supplementary figure S10**) to derive K_d_ values (**table 1, supplementary figure S8b**). The FP assay shows IRF4 mutants K59N, R98A and R98A/C99A to be compromised in their affinity for DNA compared to wild type, and all other mutants to bind DNA with greater affinity than wild type IRF4, contrary to our initial hypothesis.

To corroborate the observations made in the FP assay and attempt to elucidate the stoichiometry of interaction, we used surface plasmon resonance to determine the K_D_ of the protein-DNA interactions of the wild-type and mutant IRF4 DBDs. For this work, a 3’-biotin tag was used and the resulting biotinylated IRE hairpin was captured on a streptavidin surface. As a negative control, a hairpin with a scrambled sequence (scrIRE) of equal molecular weight and GC content was designed (scrIRE sequence with TATA to induce hairpin: 5’-AAGCATACAGAGAGACGCATATATGCGTCTCTCTGTATGCTT-3’). Therefore, measurements were referenced to a control surface of either streptavidin or treptavidin-captured biotinylated scrIRE hairpin (**supplementary figure S11**). Protein was titrated against immobilised DNA at concentrations ranging from 0.1 – 4500 nM (**figure 3a**).

**Figure 3.**
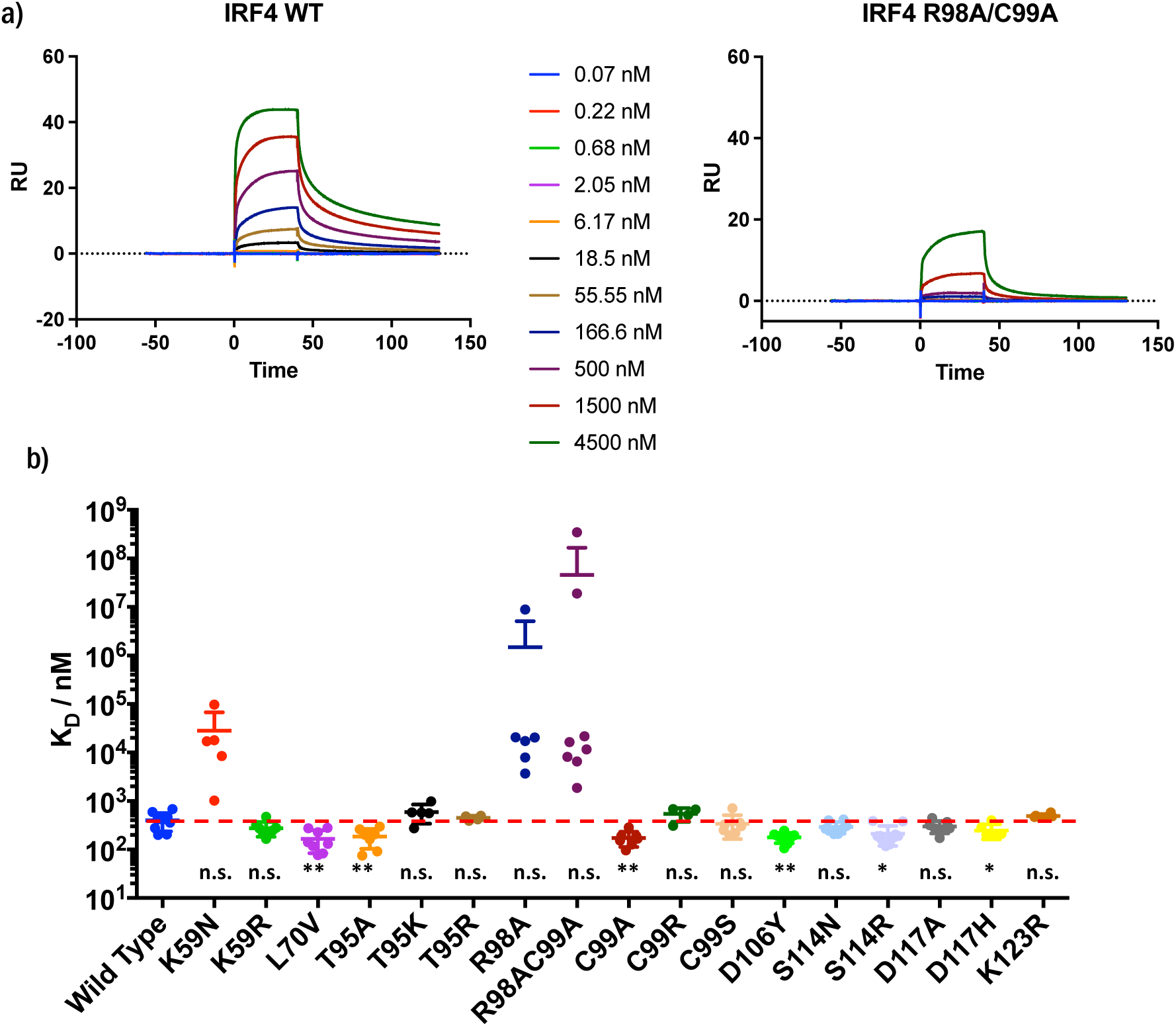
**(a)** DNA-binding sensorgrams for the wild-type IRF4 and the binding-ablated mutants R98A and R98A/C99A. Exemplar sensorgrams are presented in figure S10. **(b)** DNA-binding affinity of each disease-related mutant as determined by SPR, fit to a steady-state binding model and presented as scatter plot with mean and standard deviation.

The K_D_ values represented in **figure 3b** were calculated by fitting the binding levels (Rmax), plotted against protein concentration, to a 1:1 steady state affinity model from data across two independently prepared protein batches (**supplementary figure S13a**) and four independently prepared chip surfaces (**supplementary figure S13b**). Exemplar sensorgrams for wild-type and mutant IRF4 DBDs, with referencing to either a blank reference surface (2–1) or surface derivatised with scrIRE (4–3), are shown in **supplementary figure S11**.

While we see equivalent signals when comparing binding to either a scrambled or non-DNA control, we observe varying Rmax values over the cohort of IRF4 DBDs exceeding a theoretical 1:1 binding signal, suggesting a second weak binding event despite only one canonical IRE binding motif being present in our DNA hairpin. Assuming all DNA immobilised to the chip surface (41-58 RU) is fully functional and available for binding, we would anticipate average Rmax values of 41-58 RU for a 1:1 binding event, and 20-29 RU for a 2:1 binding event (expected Rmax = (immobilised DNA RU * immobilised DNA molecular weight) / (analyte protein molecular weight * stoichiometry)). We observed that statistically better fitting data (as determined by Chi2 value) were obtained for all sensorgrams using a two-site model (**table S3**) rather than a one-state model (**table S4**), and moreover the fitted Rmax values better represent the experimental data. The two-state binding model is therefore potentially capturing some information about non-specific/weak DNA binding to the PU.1 sequence in the fitting of a secondary binding event, as supported by the *kd2* quantified in the molar range (**supplementary figure S13c**). The affinities (K_D_) derived from the steady-state analysis and the kinetic parameters derived from the two-state kinetic fits are therefore presented for discussion (**table 1**). To investigate this more deeply, we prepared exemplary sensorgrams for each of the IRF4 samples in which we subtracted the blank reference surface data from the scrambled DNA surface to derive a measure of the binding of the proteins to the non-cognate DNA (3-1**, supplementary figure S14**). By this measure, all mutants show weak (C99R, max response ∼5 RU), or negligible non-specific binding, suggesting the second binding event observed from the two-state model is weakly specific and measurable in SPR due to local high concentrations of protein.

Finally, we used molecular dynamics simulations and the MM/PBSA method, which uses molecular mechanics combined with Poisson-Boltzmann and surface area solvation, to calculate binding energies and substantiate our observations. Typically used for ranking molecular docking outputs, MM/PBSA has been demonstrated as a useful tool for the investigation of protein-nucleic acid interactions *in silico*.(52, 53) Key to deriving useful and informative data are replicate, converged simulations of the protein-DNA complex. To that end, we simulated in triplicate the wild-type IRF4 and each of the cancer-associated mutants in complex with the IRE DNA as observed in the DNA/IRF4/PU.1 ternary complex (PU.1 removed).(15) MM/PBSA energies were extracted for each individual simulation and compared to the average of the three wild-type measurements. The results of this analysis, and comparison to the experimental two-state kinetic affinities for *kd1*, are shown in **supplementary figure S12**. In agreement with the biophysical data, although the method is not sensitive enough to differentiate wild type IRF4 and mutants that are capable of binding DNA, the mutants R98A and R98A/C99A are clearly delineated from the cohort as weaker interactors.

### Mutation of R98A ablates DNA-binding through abolition of key hydrogen bonds

As previously shown, (18) mutation of R98A in the presence or absence of an additional C99A mutation completely ablates DNA-binding (**table 1**) compared to the wild-type IRF4. Interestingly, while the R98A and C99A mutations individually have no effect on protein melting temperature (**supplementary figure S8a**), the R98A/C99A double mutation confers a significant increase in protein thermostability of mean ΔTm +1.7°C. Compared to WT, the single C99A mutant shows a 2.3-fold increase in affinity for DNA as measured by either FP or SPR (**supplementary figure S13d**). We observe a wide spread of data points in the SPR (**figure S13a**) and FP (**figure 3b**) assay results for the R98A, and R98A/C99A mutants, which is not attributable to protein batch or chip identity and could represent the difficulty in fitting binding constants where the highest concentrations in the assay are significantly less than the binding affinity.

Analysis of the kinetic fitting to the SPR data indicates the C99A mutant has a slightly increased association rate but similar dissociation rate to the wild-type IRF4 (**supplementary figure S13e-g**); the R98A and R98A/C99A mutants are notably slower *on* and faster *off*. This decrease in on-rate for R98A-containing mutants is as expected since this residue is responsible for base-specific hydrogen bonding interactions. This is recapitulated by the MM/PBSA calculations on molecular dynamics simulation data of the protein-DNA complexes, which clearly delineate the R98A and R98A/C99A mutants from the rest of the mutant and wild-type cohort (**supplementary figure S12**). Ultimately these results confirm the crucial role of specific side-chain:base interactions in IRF4 DNA-binding recognition and stability.

In contrast, we prepared the IRF4 DBD mutants D117A and D117H which had been demonstrated by Yang *et al.*(30) to be non-toxic, functional DNA-binders; the D117 mutants D117A and D117H are fully competent in DNA-binding, with enhancement observed in the FP assay (5- and 4-fold over wild-type IRF4 respectively). Residue D117 is not involved in DNA-interaction, however, mutations to this loop have been seen to reduce ternary complex formation by up to 90%(30). D117 mutations may therefore play a role in promotion of homodimerization over, for example, SPIB heterodimerisation. Enhanced DNA-binding was also observed for the CLL-related mutation L116R adjacent to D117.(13)

### Cancer-associated mutants remain competent DNA-binders

The recombinantly expressed IRF4 DBDs bearing cancer-associated mutants were compared to wild-type and the results shown in **table 1** demonstrate that, despite small significant changes in thermal melting point, all proteins were folded at physiological temperatures, and many mutants show *enhanced* DNA-binding affinity by either or both biophysical methods employed. The homogenous FP assay shows this enhancement more clearly than the surface-based SPR, perhaps owing to the increased conformational freedom for both protein and DNA in solution. The full results with statistical significance are presented graphically in **figure 3b** (SPR) **supplementary figures S8** (FP), with fold-changes compared in **supplementary figure S13d**.

One exception to the enhancement effect is K59N, which demonstrates a 10-fold decrease in average binding affinity by FP and a 70-fold decrease by SPR, though the spread of the data is wide (**supplementary figure S13d**). In stark comparison, the K59R mutant shows a slight, non-significant increase in DNA-binding affinity by 1.5-fold in both assays. By both FP and SPR, we observe strong, statistically significant increases in DNA-binding affinity for L70V of 2 to 4-fold. The T95A mutant also shows a 2-to-5-fold increase in affinity, whilst the T95R mutant demonstrates a 3.7-fold increase in affinity by FP but is essentially unchanged when compared to wild-type by SPR. The T95K mutant shows up to 2-fold increase in average affinity for DNA by FP (though 1.5-fold decreased by SPR). The C99R mutant demonstrates no significant change in binding affinity for DNA by SPR, whilst the 2-fold increase in affinity measured by FP is not statistically significant. The C99S mutation, introduced for crystallisation, shows a slight increase in DNA-binding affinity by FP (2-fold), though not by SPR. The D106Y mutant has significantly increased DNA-binding affinity: 2-fold as measured in both FP and SPR; the K123R mutant in the homogenous FP assay format (though not SPR) shows a significant increase in affinity (4-fold) over wild-type. The residue S114 lies on the same flexible loop as D117, our positive control mutant. By SPR these mutants show limited (within 2-fold) increases in DNA-binding affinity, and only S114N shows a notable 3-fold increase in binding affinity, by FP.

Finally, we inspected the k_on_ and k_off_ rates (**supplementary figures S13e-g**), obtained via the fitting of the two-state kinetic binding model to the SPR sensorgrams, for any differences between the cancer-associated mutants and the wild-type protein. Interestingly, all cancer-associated mutants demonstrate faster k_on_ rates than wild-type, suggesting the cancer-associated mutants are quicker to associate with DNA. The most notably affected are L70V, T95K, and D106Y, which are all more than 10-fold faster in their association rate. Except for S114R, all cancer-associated mutants also demonstrate faster k_off_ rates (**supplementary table S4**). The resulting K_D_s from this analysis are tabulated in **table 1** and show good agreement with the data obtained by fitting a steady-state affinity.

### Mutant K123R IRF4 possesses higher transcriptional activity relative to WT

The mutations K123R and K59R were further investigated for their ability to enhance transcription, selected as exemplar MM and TCL mutants respectively, with R98A/C99A included as a negative control. HEK293T were co-transfected with plasmid encoding full length IRF4 WT, K59R, or K123R, a reporter Firefly Luciferase (FireflyL) plasmid containing an IRF4-regulated promoter sequence, and a Renilla Luciferase transfection control. Empty vector GFP and firefly values were used for background correction. Relative to WT, the R98A/C99A double mutant IRF4 is unable to induce luciferase expression (***, p=<0.001, n=3), and a modest 1.2-fold induction of FireflyL (*, p=0.039, n=3) was demonstrated by the TCL mutant K59R (**figure 4a**), whereas K123R resulted in a significant 4.2-fold induction of FireflyL (**, p<0.01, n=3).

**Figure 4.**
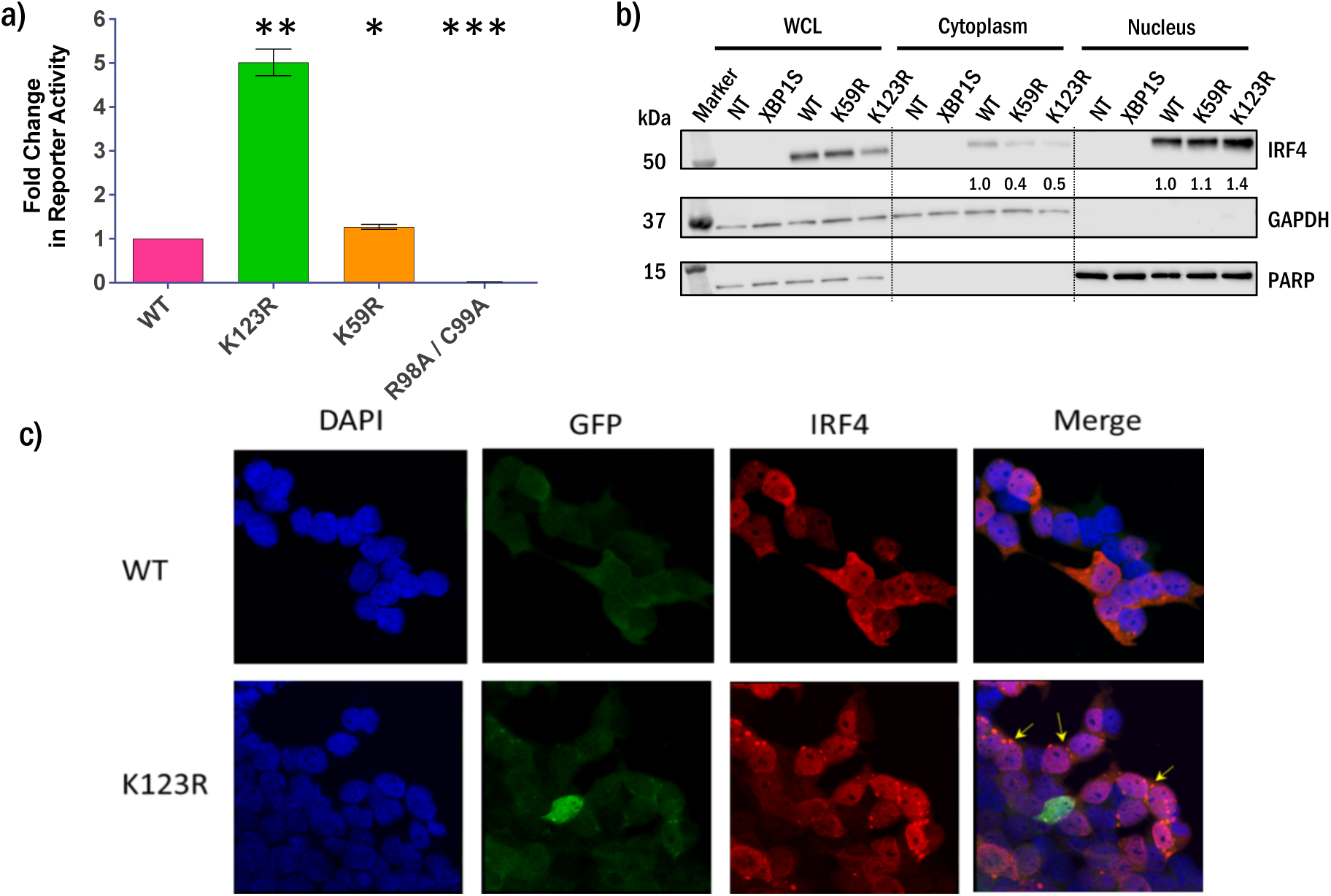
(**a**) Fold change in GFP reporter induction compared to WT for IRF4 K59R, K123R and R98A/C99A. Results of five biological replicates expressed as mean ± standard error of the mean (SEM). Significance was calculated using an unpaired student’s t-test, **(p<0.01), n=5. **(b)** HEK293T transfected with XBP1S, WT, K59R or K123R were extracted as whole cell lysates or subcellular fractions and subject to immunoblotting for IRF4. The absence of GAPDH in the nuclear fraction and PARP in the cytoplasmic fraction shows purity of the fractions. Samples were normalised to GAPDH or PARP and quantified, with numbers in bold depicting fold-change compared to WT. Images are representative of three biological repeats. NT, non-treated cells. (**c**) IF demonstrates differential subcellular localisation of K123R mutant IRF4. The nucleus is DAPI stained in blue, GFP is stained in green and IRF4 in red, with image overlay (merge) depicted to the right. Arrows depict cytoplasmic aggregates.

### K123R mutation decreases cytosolic IRF4 expression while increasing nuclear expression

To determine whether K123R or K59R mutation influences the subcellular localisation of IRF4, HEK293T were transfected with plasmid encoding XBP1S, or IRF4 WT, K123R or K59R. After 24 h, whole cell lysate and nuclear and cytoplasmic fractions were extracted, and protein was separated by SDS-PAGE and subject to immunoblotting. Loading normalisation was by comparison to GAPDH for whole cell lysates and cytoplasmic fractions, and PARP for nuclear fractions (**figure 4b**). Compared to WT, total IRF4 expression is 0.5- and 0.7-fold lower in K123R and K59R-transfected cells, respectively. Cytosolic IRF4 expression is 0.5-fold lower in K123R and 0.4-fold lower in K59R transfected cells, compared to WT. Nuclear expression of IRF4 is increased 1.4-fold in K123R and 1.1-fold in K59R transfected cells, relative to WT. K123R IRF4 therefore has lower cytoplasmic expression and increased nuclear expression compared to WT (verified by 3 technical replicates of 3 biological replicates, representative example shown).

### Differential subcellular localisation of WT and K123R IRF4 by immunofluorescence

HEK293T were seeded onto coverslips, transfected with pIRES2.EGFP plasmids, stained with antibodies, and visualised by microscopy to determine the subcellular localisation of the IRF4 protein (**figure 4c**). K123R mutant IRF4 demonstrated increased nuclear expression compared to WT, as seen by a stronger merged signal from co-localisation of DAPI and IRF4 (pink). K123R IRF4 also had less intense IRF4 staining in the cytoplasm relative to WT. This result was confirmed by three independent biological repeats. It is noted that cytoplasmic aggregates containing IRF4 are present, but it is believed these are the consequence of cell processing of the exogenous protein and intense staining as trafficking through the Golgi in the perinuclear space occurs. Protein expression effects were investigated using a range of plasmid doses from 0.5 µg to 1.0 µg; at 0.5 µg, higher nuclear localisation and lower cytoplasmic expression of mutant IRF4 compared to WT was still detected and at 1.0 µg this effect was more pronounced, indicating a plasmid concentration-dependent effect on expression (**supplementary figure S15**). Notably, focal aggregates are present at higher concentrations of IRF4, suggesting these are caused by overexpression.

## Discussion

The mutational landscape of IRF4 in MM and TCL indicated that DBD mutations may confer better prognosis, which we hypothesised could reflect differential DNA-binding. Weaker DNA-binding of IRF4 would be expected to reduce IRF4-addicted signalling, potentially influencing the outcome to treatment. However, the better prognosis of MM with mutated IRF4 following triple immunotherapy drug treatment (either cyclophosphamide, thalidomide, and dexamethasone, or cyclophosphamide, lenalidomide, and dexamethasone)(8), could conceivably also result from enhanced DNA-binding and a stronger IRF4 signalling drive that is more responsive to IRF4-directed therapy. Thalidomide and lenalidomide are known to down-regulate IRF4-expression specifically through a mechanism that involves drug binding to Cereblon (CRBN) and modulation of the specificity of the CRBN-CRL4 E3-ligase complex, leading to ubiquitination and degradation of the upstream transcription factors Ikaros (IKZF1) and Aiolos (IKZF3) that negatively regulate IRF4.(54) Given the significant interest in the role of IRF4 in haematological malignancies, we undertook to characterise the relative DNA-binding characteristics of a cohort of clinically relevant IRF4 DBD mutants.

To confirm the structure of the apo human DBD, we crystallised IRF4 (20–132) bearing three surface entropy mutations and a C99S mutation to prevent protein crosslinking. The surface entropy mutations facilitate crystal packing by replacing E45, E46, and K47 with alanine residues which allow neighbouring molecules to array. As demonstrated by FP and SPR, the C99S mutation has little to no effect on DNA-binding affinity. We observe that the neighbouring residue, R98, adopts a conformation seen in several of the NMR models of apo IRF4 in solution. The conservation of R98 across the IRF family indicates the base-specific interaction observed for this residue in the complex of IRF4 with DNA is crucial to its function as a transcription factor. The C99 residue is similarly conserved in all but IRF3 where it is present as a serine. Here, we utilised single C99 mutants C99A, C99S, and C99R to investigate the role of C99 in the DNA interaction, however, it is clear that R98 is driving the binding event and the adjacent residue can tolerate some change around charge, polarity and bulk.

As oncogenic mutations in the p53 DBD have been shown to reduce domain stability and induce changes in folding which in turn affect function, (55) we utilised a protein thermostability assay to investigate the effect of the IRF4 cancer-associated mutations on the stability of the recombinantly expressed DBDs. We hypothesised that a subset of the cancer-associated mutations could compromise structural integrity and so impact DNA-binding capability. In particular the L70V mutation would disrupt the hydrophobic core of the domain by introducing a smaller side-chain, thus leading to diminished functional competence. Approximately half of the tested mutants showed no significant difference in mean Tm compared to wild-type IRF4, and the R98A/C99A mutation is seen to be stabilising rather than destabilising. We confirmed, however, that the L70V mutant does indeed destabilise the DBD significantly, with a mean decrease in Tm of 4.9 °C. Smaller, yet still significant, mean decreases in Tm of 3.7 °C and 3.8 °C were measured for IRF4 mutants S114N and S114R, respectively. S114 lies on the surface of the DBD and participates in a hydrogen bond network which mediates interactions with DNA and PU.1. Though we would expect the L70V, S114N and S114R mutants to be folded at physiological temperatures, the observed decreases in Tm are perhaps indicative of an increase in protein disorder which could impact DNA- and co-factor binding.

To quantitatively characterise DNA-binding of the wild type and disease-related mutants, we utilised two methods: a fluorescence polarisation (FP) assay (**supplementary figure S8b**) to measure affinity in solution, and a surface plasmon resonance (SPR) direct binding assay to characterise the affinity and kinetics of binding to immobilised DNA (**figure 3b**). The DNA binding of the wild-type IRF4 DBD has been monitored using a variety of assay formats, namely electrophoretic mobility shift (EMSA), quantitative hydroxyl radical foot-printing (QHRF) and FP. Reported values for untagged protein establish a K_d_ in the range of 500-900 nM.(13, 56–58) The affinities measured herein by FP (565 ± 191) and SPR (402 ± 165 nM) lie within this range and are in good agreement with each other. In further agreement with the literature, the DNA-binding ability of the engineered mutants R98A and R98A/C99A has been ablated. All other mutants that we tested remained capable of binding DNA, including our positive control mutants D117A and D117H.

We analysed our SPR data for evidence of non-specific binding by comparing the blank reference surface to that functionalised with a scrambled IRE sequence (**supplementary figure S14**). This showed very little evidence of non-specific binding, with one notable exception: the C99R mutant. The C99R mutant was recently studied in the lymphoma context, particularly classic Hodgkin lymphoma.(14) In this context, the C99R mutant was observed to no longer bind the canonical ISRE (in comparison to WT IFR4) and gain function up-regulating non-canonical genes. Together with our data, we hypothesise that the introduction of an arginine, capable of base-pair interactions like the adjacent R98, may expand the range of potential protein-DNA interactions and thus alter base specificity.

It is possible that the small, but significant, changes in binding affinity that we measured have physiological relevance in a cancer setting. We therefore explored other parameters that may potentially explain the cancer-association of these hotspot mutations. First, we considered the residence time of the proteins on DNA, as measured in our SPR assay. Analysis of the kinetic fitting to the SPR data shows there are changes in association and dissociation rates across the mutant cohort. The association (k_on_**, supplementary figure S13e**) of wild-type IRF4 was measured to average in the order of 10^5^ M^-1^ s^-1^, slower than some observed protein:DNA interactions (limited to 10^8^ M^-1^ s^-1^(59)) indicative of slide-searching by IRF4 along the DNA for the optimal binding position in addition to diffusion through the flow cell.(59)

The IRF4 WT dissociation rate (k_off_**, supplementary figure S13f**) was measured in the order of 0.1 s^-1^. This off-rate is similar to that reported for other transcription factors, such as FOXO4, (60) GATA, and ER DBDs, (61) and corresponds to a residence time on DNA of approximately 5 s. The cancer-associated mutants tend to dissociate from DNA 0.2 to 2-log fold faster than the wild-type IRF4 DBD (**supplementary figure S13g**). The sensorgram curves for the binding-ablated mutants R98A and R98A/C99A can be analysed and fit to a two-state binding model; values obtained demonstrate a statistically significantly slower k_on_ and faster k_off_. The majority of cancer-associated mutants demonstrate faster k_on_ and k_off_ rates than the wild-type protein. Residence time (!**, supplementary table S4**) can be calculated as 1/k_off_, by which we observe that only S114R and C99A have residence times comparable to wild-type, with all other mutants reduced - by up to 93% in the case of T95K (0.36 s). This effect(61) has been seen in other disease-related transcription factor mutants such as STAT1, a co-factor of IRF4, where DNA-binding domain and linker mutants possessed similar binding affinities but increased dissociation rates, reducing residence time and co-factor recruitment activity.(62) In the context of full-length IRF4 as well as in the presence of co-factors, these off rates could be further modulated; however, the clear reduction in residence time for recombinant DBDs shows the essential interaction of protein and DNA is notably perturbed.

Second, we considered the effect these mutations could have on biological stability or signalling. For example, the residue S114 in IRF4 corresponds to S97 in IRF3, which is seen to be phosphorylated by TBK1.(33) It is possible that mutation of S114 not only alters the binding partner specificity of IRF4, but also interrupts phosphorylation-mediated controls. Having demonstrated through thermal melt assays their negligible impact on protein thermostability, we considered the loss of K59 and K123 may abolish ubiquitination sites or impact transactivation. Multiple E3 ligases have been identified as ubiquitylators of IRF family members, for example MDM2 for IRF2, (63) TRIM21 for IRF5, (64) TRAF6 for IRF7, (65) and Cbl for IRF8.(66) The residue K123R is equivalent to K108 in IRF8, which is a site of ubiquitination by Cbl;(66) the IRF8 K108E mutation not only causes a loss of nuclear localisation, but also transactivation. Though the K59 residue is highly conserved as a lysine or arginine residue, it does not correspond to any known site of ubiquitination in other IRF family members. However, a recent study by Cherian *et al*.(67) showed K59R in IRF4 to result in enhanced nuclear expression and speculate it may be an ablated ubiquitination site. If K59 and/or K123 are sites of IRF4 ubiquitination, then the K59R and K123R mutants may not be properly targeted for proteasomal degradation and adequately deactivated, thus enhancing IRF4 addiction. We investigated the impact of mutation on the stability of the protein upon inhibition of both translation (cycloheximide treatment) and degradation (inhibition of the proteasome by MG132) and did not observe an alteration in stability that would account for the enhanced IRF4 addiction reported for these mutations, however this may be cell type dependent (data not shown).

In order to determine whether the K123R mutation conferred additional function to IRF4, we conducted reporter assays following transfection using a construct to assess a the regulation of IRF responsive elements adjacent to the transcription start site of *ZBTB32*(22). After normalisation of expression to that of GFP from the internal ribosome entry site and correlation of the reporter activity to Renilla luciferase, we observed that K123R expression resulted in significantly greater reporter activity; luciferase activity driven from an IRF responsive element was measured at 4-fold that for WT IRF4 (p<0.01) (**figure 4a**). As we had not expressed any additional cofactors and the IRF4 function was enhanced over empty vector controls for WT IRF4, we conclude that the mutation enhances transcriptional output and that this is likely to a result of favourable interaction of the K123R mutant with DNA.

Third, we considered whether these cancer-associated mutations could affect nuclear localisation or transport. Given the previously reported enhanced nuclear localisation of K59R and the inferred impact of K123R from the multiple myeloma-associated frequency of mutation, we investigated the role of the two mutations in nuclear localisation and transcriptional activity. By analysis of the western blot data for K123R, K59R and WT IRF4 localisation, overall expression of protein (whole cell lysate), is reduced for K123R relative to WT and slightly lower in K59R (**figure 4b**). This may reflect the greater proportion of protein in the nuclear extract and the incomplete extraction of cellular protein by the whole cell lysis or may be that the production of protein by the mutant containing plasmid is lower than WT; it is not clear that the mutation *per se* results in such an effect. We do observe that both K59R and K123R mutants result in a greater ratio of nuclear localisation of expressed protein when compared to the distribution of the WT protein, in contrast with the previous report of Cherian *et al*.(67). It should be noted that conducting the experiments in HEK293 cells may not have presented the full complement of ancillary proteins and thus differential effects are not unexpected when compared to proteins expressed in the context of a haematopoietic/lymphoid cell. We confirmed the preferential nuclear localisation of the K123R IRF4 protein by immunofluorescence (**figure 4c**), wherein the expression of the mutant IRF4 is almost entirely nuclear (near complete overlay with DAPI-stained nuclei). In contrast, the WT IRF4 resulted in clear cytoplasmic staining and less pronounced nuclear localisation.

The K94-T95 residue pair in IRF4 corresponds to the K77-R78 pair in IRF3 which was seen to be essential for nuclear localisation of IRF3.(31) Moreover, K94 corresponds to K78 in IRF1, a known site of ubiquitination.(68) As the T95 mutations do not share the clinical relevance of the K123 or K59 mutations, we did not investigate their cellular role but we can therefore speculate that the T95 mutants may have a role in mediating either nuclear localisation or E3 ligase recognition, which would have the effect of enhancing or restricting signalling respectively.

It remains to be seen whether these IRF4 cancer-associated mutants are passenger mutations or drivers of disease. However, we have demonstrated small, but significant changes in quantitative parameters of DNA-binding. In the context of previous literature(13, 67) these mutations may have physiological relevance as subtle activators in TCL and MM, evidenced by the increased transcriptional activity and greater nuclear localisation of K123R IRF4. It is of interest that the K123R hotspot mutation is reported to have a favourable prognosis; our data suggest added fitness through enhanced nuclear localisation and reporter output that would favour selection of the mutation. Favourable prognosis resulting from such an activating mutation could feasibly occur through a greater responsiveness to direct IRF4 modulating therapy (i.e. thalidomide or lenalidomide treatment), or the expression of neo-antigens by K123R-mutant clones resulting in increased immunogenicity.(69)

### Conclusion

The mutational landscape of IRF4 is of great interest not only for its implications for cancer biology but also in other emergent disease settings.(70) Based on structural analysis combined with mutation prediction software, we hypothesised that cancer-associated mutations in the IRF4 DNA-binding domain were likely to destabilise and disfavour protein-DNA interactions. Our crystal structure of an IRF4 DBD surface entropy mutant in the apo state supported this view with the protein alone and within the context of DNA. We aimed to examine IRF4 DBD mutants experimentally, also utilising two literature-precedent control mutants known to ablate DNA-binding: the R98A and R98A/C99A double mutant, through which we confirmed the essential role of the R98 residue in DNA-binding.

We characterised the changes in protein integrity resulting from introduced mutations by thermal melt assays, demonstrating that while seven out of 17 mutations reduced the Tm, the resultant Tm’s remained well above physiological temperature and are therefore unlikely to be responsible for loss of function. Subsequent analysis of binding to DNA demonstrated that 15 out of 17 mutants bound DNA with affinities similar to or tighter than wild-type, whether assayed in a homogenous assay format (fluorescence polarisation) or utilising a surface-immobilised method (surface plasmon resonance). Further analysis of the kinetic binding profiles showed that the cancer-associated IRF4 DBD mutants bind DNA faster than wild-type, but also dissociate more rapidly, with some k_off_ rates increasing by up to two-log fold and approaching the limit of detection by the SPR instrument.

Collectively, these results indicate that cancer-associated mutations in the IRF4 DBD confer changes which act to modestly reduce thermostability in the majority (7/17) of cases, and increase the dissociation rate from DNA, whilst maintaining functional DNA-binding ability of similar or tighter binding affinity. Preliminary biological investigation of two clinically relevant mutations, K123R and K59R, show the K123R mutation confers a significantly higher transcriptional activity than WT, and both mutants show distribution skewed toward nuclear residence. We conclude that whatever the role of the mutation beyond stability and DNA-binding, the enhanced nuclear localisation and transcriptional activity of K123R IRF4 support the notion that it is an activating mutation that can be preferentially selected for in multiple myeloma.

## Data Availability

All raw data are available on request to the corresponding author.

## Acknowledgements

The authors would like to acknowledge Arnaud Basle, for provision of crystallisation facility and in-house X-ray generator access; Tom Davies, for assistance with diffraction data analysis; Alex Dias, beamline scientist and data collection at DIAMOND; Tim Kirk for software and crystallographic computing support; Jane Endicott, for input to primer design; Richard Heath, for provision of modified vectors for expression; Finn Holding, for initial LC-MS data; Judith Reeks, for preliminary FP assay development; Pamela Williams for assistance with data collection and many helpful discussions regarding optimisation of crystals; and Sandra Grieve for helpful discussions. The authors would like to thank Diamond Light Source for beamtime (experiment number in-15437-3). For the purpose of open access, the authors have applied a Creative Commons Attribution (CC BY) licence to any Author Accepted Manuscript version arising from this submission.

