## Supplemental Material for "Structural, biophysical and biological analysis and characterisation of IRF4 DNA-binding domain mutations associated with multiple myeloma"

**Tatum, Scott, *et al*. (2020) Supplementary Information**


| | | |
|---|---|---|
| Table S1 | Tabulated summary of mutation prediction software output. | 2 |
| Table S2 | Tabulated observed masses of IRF4 proteins following intact protein LC-MS analysis. | 4 |
| Table S3 | Crystallographic data processing and refinement statistics. | 5 |
| Table S4 | Experimental average $K_D$s ± SD derived by fitting of two-state kinetic binding model to SPR data. | 6 |
| Table S5 | Experimental average $K_D$s ± SD derived by fitting of one-state kinetic binding model to SPR data. | 7 |
| Table S6 | Forward and reverse mutagenesis primers. Mutation sites are underlined in the forward primer. | 8 |
| Note S1 | Expression tag sequences. | 9 |
| Figure S1 | Interactions made by K59 | 10 |
| Figure S2 | Interactions made by T95 | 11 |
| Figure S3 | Interactions made by D106 | 12 |
| Figure S4 | Interactions made by S114 | 13 |
| Figure S5 | Interactions made by K123 | 14 |
| Figure S6 | Interactions made by L70 | 15 |
| Figure S7 | Interactions made by C99 | 16 |
| Figure S8 | Graphed thermal melt data and results of FP assay for IRF4 WT and all mutants. | 17 |
| Figure S9 | Exemplar LCMS raw data and deconvoluted mass spectra for IRF4 WT, K59R, K123R. | 18 |
| Figure S10 | Exemplar fluorescence polarisation curves for IRF4 WT and all mutants. | 19 |
| Figure S11 | Exemplar SPR sensorgrams for IRF4 WT and all mutants. | 20 |
| Figure S12 | Comparison of calculated binding energy and experimental binding affinity for IRF4 WT and all mutants. | 22 |
| Figure S13 | Representations of SPR data for IRF4 WT and all mutants – (a) protein batch variation, (b) chip surface variation, (c) one *vs.* two-state binding models, (d) fold change in affinity by SPR and FP, (e) on-rates, (f) off-rates, (g) log fold change in on/off rates. | 23 |
| Figure S14 | Sensorgrams for binding to scrambled control DNA demonstrating lack of non-specific binding for IRF4 WT and all mutants.. | 25 |
| Figure S15 | Immunofluorescence demonstrating subcellular localisation of IRF4 WT, K59R and K123R. | 27 |

**Table S1:** Tabulated summary of mutation prediction software output.

| | **Mutation Taster**(28) † | | | **PROVEAN**(29)‡ | | **SIFT**(30)¥ | | **CADD**(31)§ | |
|---|---|---|---|---|---|---|---|---|---|
| **Variant** | **Prediction** | **P** | **Splicing** | **Delta Score** | **Prediction** | **Prediction** | **Score** | **Raw Score** | **PHRED** |
| I49V | disease causing | 1.00 | Alteration | -0.507 | Neutral | Tolerated | 0.24 | 2.137 | 20.7 |
| K59N | disease causing | 1.00 | None | -4.434 | Deleterious | Predicted not tolerated | not given | not annotated | n/a |
| K59R | disease causing | 1.00 | Alteration | -2.528 | Deleterious | Tolerated | 0.72 | 4.192 | 31 |
| L70V | disease causing | 1.00 | Alteration | -1.805 | Neutral | Tolerated | 0.09 | 4.444 | 33 |
| T95A | disease causing | 1.00 | Alteration | -4.57 | Deleterious | Alignment suggests tolerated | not given | not annotated | n/a |
| T95K | disease causing | 1.00 | Alteration | -5.644 | Deleterious | Predicted not tolerated | not given | 3.614 | 25.5 |
| T95R | disease causing | 1.00 | Alteration | -5.644 | Deleterious | Affect protein function | 0.02 | 3.492 | 25.1 |
| R98A | disease causing | 1.00 | None | -9.636 | Deleterious | Affect protein function | 0.00 | not annotated | n/a |
| C99S | disease causing | 1.00 | Alteration | -8.673 | Deleterious | Predicted not tolerated | not given | not annotated | n/a |
| C99A | disease causing | 1.00 | None | -11.563 | Deleterious | Predicted not tolerated | not given | not annotated | n/a |
| C99R | disease causing | 1.00 | Alteration | -5.735 | Deleterious | Affect protein function | 0.00 | 3.293 | 24.4 |
| D106Y | disease causing | 1.00 | Alteration | -8.135 | Deleterious | Affect protein function | 0.00 | 4.288 | 32 |
| S114N | disease causing | 1.00 | Alteration | -2.236 | Neutral | Predicted tolerated | not given | 4.143 | 29.6 |
| S114R | disease causing | 1.00 | None | -4.336 | Deleterious | Tolerated | 0.07 | 3.544 | 25.3 |
| D117A | disease causing | 1.00 | Alteration | -7.104 | Deleterious | Predicted tolerated | not given | 4.267 | 32 |
| D117H | disease causing | 1.00 | Alteration | -6.152 | Deleterious | Tolerated | 0.11 | 4.343 | 32 |
| K123R | disease causing | 1.00 | Alteration | -2.896 | Deleterious | Affect protein function* | 0.00 | 3.590 | 25.4 |

†**MutationTaster** predicts alterations as either disease causing (probably deleterious), disease causing automatic (meaning it is known to be deleterious), polymorphism (probably harmless), or polymorphism automatic (known to be harmless). The probability value represents the 'security' of the prediction, and splice site predictions are derived via NNSplice within MutationTaster. ‡**PROVEAN** calculates a delta score based on pairwise sequence alignments; a negative score indicates a deleterious effect whereas a positive score can be interpreted as neutral. For this analysis, PROVEAN used 30 clusters with 173 supporting sequences. ¥**SIFT** predicts the effect of coding variants based on sequence homology. The SIFT score is a normalised probability where a value from 0.00 - 0.05 is predicted to

affect protein function. (* - SIFT warned the sequences used were not diverse enough to give full confidence in this prediction.) [§]**CADD** gives a C score derived from a support vector machine trained to measure deleteriousness. The raw C score can be interpreted as the annotation profile where higher relative scores represent increased likelihood the variant has a deleterious effect. The PHRED score is a scaled C score from 1 - 99 which represents the rank in order of magnitude (*i.e.* variants in the top 10% of deleterious mutations are scored >10, variants in the top 1% of deleterious mutations are >20, top 0.1% are >30 etc.). Where no SNV is present in COSMIC, no CADD analysis was performed.

Table S2: Observed masses of IRF4 proteins following intact protein LC-MS analysis

| Protein | Calculated Mass (Da) | Observed mass (Da) | Error (Da) |
|---|---|---|---|
| Wild-type | 13455.3 | 13455.3 | 0.0 |
| K59N | 13441.2 | 13441.0 | -0.2 |
| K59R | 13483.3 | 13482.3 | -1.0 |
| L70V | 13441.3 | 13440.5 | -0.8 |
| T95A | 13425.3 | 13424.9 | -0.4 |
| T95K | 13482.3 | 13481.6 | -0.6 |
| T95R | 12510.3 | 13509.2 | -1.1 |
| R98A | 13370.2 | 13369.9 | -0.3 |
| R98A/C99A | 13338.1 | 13338.5 | 0.4 |
| C99A | 13423.1 | 13423.6 | 0.5 |
| C99R | 13508.3 | 13507.8 | -0.5 |
| C99S | 13439.2 | 13439.7 | 0.5 |
| D106Y | 13503.4 | 13502.9 | -0.5 |
| S114N | 13482.3 | 13483.0 | 0.7 |
| S114R | 13524.4 | 13524.0 | -0.4 |
| D117A | 13411.2 | 13410.5 | -0.7 |
| D117H | 13477.2 | 13477.1 | -0.1 |
| K123R | 13483.3 | 13482.8 | -0.5 |

**Table S3:** Crystallographic data processing and refinement statistics

| | |
|---|---|
| PDB ID | 6TD4 |
| Beam-line | Diamond Light Source, I04-1 |
| Wavelength (Å) | 0.92 |
| Detector type | Dectris Pilatus 6M-F |
| Data collection date | 14/07/2016 |
| Space group | $C222_1$ |
| Cell constants a; b; c (Å) | 66.9, 67.5, 69.9 |
| Resolution range (Å)[1] | 39.28-1.71 (1.78-1.71) |
| Completeness overall (%)[1] | 100 (100) |
| Reflections, unique [1] | 17417 (1727) |
| Multiplicity[1] | 6.5 (5.7) |
| Mean(I)/sd(I)[1] | 16.9 (1.7) |
| $R$meas $_{overall}$ [2] | 0.041 (1.120) |
| CC½[1] | 0.994 (0.839) |
| $R$value $_{overall}$ (%) [3] | 21.65 |
| $R$value $_{free}$ (%)[1] | 24.50 |
| Non hydrogen protein atoms | 887 |
| Non hydrogen ligand atoms | 1 |
| Solvent molecules | 135 |
| R.m.s. deviations from ideal values | |
| Bond lengths (Å) | 0.004 |
| Bond angles (°) | 0.590 |
| Average *B* values (Å$^2$) | |
| Average B values for protein (Å$^2$) | 51.6 |
| Average B values for ligand (Å$^2$) | 89.3 |
| Average B values for water (Å$^2$) | 59.4 |
| Φ, Ψ angle distribution for residues [4] | |
| In favoured regions (%) | 96.1 |
| In allowed regions (%) | 3.9 |
| In outlier regions (%) | 0.0 |

**1** Values in parentheses refer to the outer resolution shell
**2** $R_{meas} = (\Sigma_{hkl} \sqrt{[n/(n-1)]}\, \Sigma_i |I_i - \langle I \rangle|) / \Sigma_{hkl} \Sigma_i I_i$
**3** $R_{value} = S_{hkl} ||F_{obs}| - |F_{calc}|| / S_{hkl} |F_{obs}|$
$R_{free}$ is the cross-validation $R$ factor computed for the test set of 5% of unique reflections
**4** Ramachandran statistics as defined by Coot

**Table S4:** Experimental average $K_D$s ± SD derived by fitting a two-state kinetic binding model to SPR data. Curves were fit individually per experiment and derived values averaged. Curves which were fit with kd rates exceeding 2 $s^{-1}$ (the upper limit of detection specified for the Biacore S200) were excluded from the analysis. Residence time (τ) was calculated as 1/kd1.

| Protein | ka1 (1/Ms) | kd1 (1/s) | ka2 (1/s) | kd2 (1/s) | $K_D$ ± SD (nM) | $R_{max}$ (RU) | Chi$^2$ | τ / s |
|---|---|---|---|---|---|---|---|---|
| Wild-type | 3.17E+05 | 1.94E-01 | 2.57E-02 | 9.07E-03 | 187 ± 143 | 35.52 | 0.32 | 5.15 |
| K59N | 5.31E+05 | 6.14E-01 | 3.00E-02 | 2.95E-03 | 107 ± 133 | 15.96 | 0.09 | 1.63 |
| K59R | 1.19E+06 | 6.97E-01 | 1.23E-02 | 1.45E-02 | 460 ± 429 | 36.15 | 0.43 | 1.43 |
| L70V | 5.79E+06 | 5.53E-01 | 7.83E-03 | 1.04E-02 | 121 ± 111 | 28.95 | 0.42 | 1.81 |
| T95A | 2.17E+06 | 1.38E+00 | 8.89E-03 | 1.24E-02 | 305 ± 174 | 34.91 | 0.65 | 0.72 |
| T95K | 3.03E+06 | 2.74E+00 | 2.29E-03 | 8.81E-03 | 913 ± 147 | 36.42 | 0.28 | 0.36 |
| T95R | 1.01E+06 | 5.20E-01 | 1.31E-03 | 8.01E-02 | 481 ± 149 | 34.12 | 0.55 | 1.92 |
| R98A | 3.38E+04 | 6.57E-01 | 7.66E-03 | 2.93E-02 | > 4500 | 104.99 | 0.02 | 1.52 |
| R98A/C99A | 4.44E+04 | 8.55E-01 | 1.46E-02 | 4.36E-02 | > 4500 | 50.91 | 0.01 | 1.17 |
| C99A | 1.14E+06 | 1.83E-01 | 1.93E-02 | 1.13E-02 | 61 ± 29 | 51.67 | 1.20 | 5.46 |
| C99R | 7.70E+05 | 5.78E-01 | 7.13E-03 | 1.29E-02 | 660 ± 317 | 82.87 | 0.68 | 1.73 |
| C99S | 1.02E+06 | 2.76E-01 | 1.75E-02 | 1.61E-02 | 139 ± 43 | 39.25 | 0.26 | 3.62 |
| D106Y | 3.98E+06 | 4.60E-01 | 8.06E-03 | 9.37E-03 | 135 ± 121 | 27.76 | 0.49 | 2.17 |
| S114N | 9.40E+05 | 3.58E-01 | 2.35E-02 | 7.68E-03 | 145 ± 151 | 45.11 | 0.40 | 2.79 |
| S114R | 8.12E+05 | 1.60E-01 | 2.02E-02 | 5.18E-03 | 49 ± 26 | 41.57 | 0.60 | 6.25 |
| D117A | 1.75E+06 | 9.20E-01 | 1.46E-02 | 3.79E-02 | 293 ± 213 | 56.67 | 0.53 | 1.09 |
| D117H | 4.19E+06 | 1.18E+00 | 1.62E-02 | 1.10E-02 | 204 ± 131 | 51.22 | 0.44 | 0.85 |
| K123R | 1.29E+06 | 1.73E+00 | 1.47E-03 | 4.53E-03 | 1107 ± 341 | 46.26 | 1.30 | 0.58 |

**Table S5:** Experimental average $K_D$s ± SD derived by fitting a one-state kinetic binding model to SPR data. Curves were fit individually per experiment and derived values averaged.

| Protein | ka (1/Ms) | kd (1/s) | $K_D$ (nM) | $R_{max}$ (RU) | $Chi^2$ |
|---|---|---|---|---|---|
| Wild-type | 1.81E+05 | 1.23E-02 | 166 ± 106 | 25.74 | 0.85 |
| K59N | 3.22E+05 | 3.76E-01 | 347 ± 685 | 9.92 | 0.12 |
| K59R | 1.49E+06 | 2.21E+01 | 147 ± 153 | 26.89 | 0.78 |
| L70V | 2.44E+06 | 8.00E-02 | 72 ± 63 | 13.91 | 0.41 |
| T95A | 9.55E+05 | 7.94E-02 | 191 ± 250 | 15.02 | 0.46 |
| T95K | 6.40E+05 | 3.33E-01 | 582 ± 257 | 26.40 | 0.52 |
| T95R | 7.04E+05 | 2.21E-01 | 336 ± 73 | 24.58 | 0.63 |
| R98A | 2.32E+04 | 2.07E-01 | > 4500 | 74.08 | 0.41 |
| R98A/C99A | 6.66E+04 | 1.40E-01 | 4339 ± 3025 | 11.80 | 0.05 |
| C99A | 3.66E+05 | 1.99E-02 | 52 ± 20 | 31.25 | 1.99 |
| C99R | 2.83E+05 | 1.01E-01 | 461 ± 358 | 56.63 | 2.44 |
| C99S | 1.13E+08 | 2.17E+01 | 140 ± 63 | 28.42 | 0.93 |
| D106Y | 2.95E+05 | 1.72E-02 | 147 ± 206 | 22.55 | 1.04 |
| S114N | 2.15E+05 | 1.92E-02 | 127 ± 103 | 27.26 | 0.87 |
| S114R | 5.17E+05 | 1.34E-02 | 47 ± 21 | 27.52 | 1.19 |
| D117A | 2.86E+05 | 1.11E-01 | 260 ± 169 | 44.62 | 1.63 |
| D117H | 5.01E+05 | 9.82E-02 | 174 ± 93 | 33.33 | 0.90 |
| K123R | 3.96E+05 | 2.88E-01 | 942 ± 374 | 20.67 | 0.42 |

**Table S6:** Forward and reverse mutagenesis primers. Mutation sites are underlined in the forward primer.

| IRF4 Mutant | Forward Primer Sequence | Reverse Primer Sequence |
|---|---|---|
| E45A E46A K47A | GCT GGT GTG GGA GAA CGC GGC CGC GAG CAT CTT CCG CAT CC | GGA TGC GGA AGA TGC TCG CGG CCG CGT TCT CCC ACA CCA GC |
| K59N | CC TGG AAG CAC GCG GGC AAC CAG GAC TAC AAC CGC GAG | CTC GCG GTT GTA GTC CTG GTT GCC CGC GTG CTT CCA GG |
| K59R | CC TGG AAG CAC GCG GGC AGA CAG GAC TAC AAC CGC GAG | CTC GCG GTT GTA GTC CTG TCT GCC CGC GTG CTT CCA GG |
| L70V | CGC GAG GAG GAC GCC GCG GTG TTC AAG GCT TGG GCA CTG TTT AAA G | C TTT AAA CAG TGC CCA AGC CTT GAA CAC CGC GGC GTC CTC CTC GCG |
| T95A | CG GAC CCT CCC ACC TGG AAG GCG CGC CTG CGG TGC GCT TTG AAC AAG AG | CT CTT GTT CAA AGC GCA CCG CAG GCG CGC CTT CCA GGT GGG AGG GTC CG |
| T95K | CG GAC CCT CCC ACC TGG AAG AAA CGC CTG CGG TGC GCT TTG AAC AAG AG | CT CTT GTT CAA AGC GCA CCG CAG GCG TTT CTT CCA GGT GGG AGG GTC CG |
| T95R | CG GAC CCT CCC ACC TGG AAG AGA CGC CTG CGG TGC GCT TTG AAC AAG AG | CT CTT GTT CAA AGC GCA CCG CAG GCG TCT CTT CCA GGT GGG AGG GTC CG |
| R98A | C TGG AAG ACG CGC CTG GCG TGC GCT TTG AAC AAG AGC | GCT CTT GTT CAA AGC GCA CGC CAG GCG CGT CTT CCA G |
| R98AC99A | C TGG AAG ACG CGC CTG GCG GCC GCT TTG AAC AAG AGC | GCT CTT GTT CAA AGC GGC CGC CAG GCG CGT CTT CCA G |
| C99A | G AAG ACG CGC CTG CGG GCC GCT TTG AAC AAG AGC | GCT CTT GTT CAA AGC GGC CCG CAG GCG CGT CTT C |
| C99R | C TGG AAG ACG CGC CTG CGG AGA GCT TTG AAC AAG AG | CT CTT GTT CAA AGC TCT CCG CAG GCG CGT CTT CCA G |
| C99S | GAA GAC GCG CCT GCG GTC CGC TTT GAA CAA GAG C | GCT CTT GTT CAA AGC GGA CCG CAG GCG CGT CTT C |
| D106Y | GCT TTG AAC AAG AGC AAT TAT TTT GAG GAA CTG GTT GAG C | G CTC AAC CAG TTC CTC AAA ATA ATT GCT CTT GTT CAA AGC |
| S114N | GAA CTG GTT GAG CGG AAC CAG CTG GAC ATC TCA GAC C | G GTC TGA GAT GTC CAG CTG GTT CCG CTC AAC CAG TTC |
| S114R | GAA CTG GTT GAG CGG AGA CAG CTG GAC ATC TCA GAC C | G GTC TGA GAT GTC CAG CTG TCT CCG CTC AAC CAG TTC |
| D117A | GTT GAG CGG AGC CAG CTG GCG ATC TCA GAC CCG TAC AAA GTG TAC AG | CT GTA CAC TTT GTA CGG GTC TGA GAT CGC CAG CTG GCT CCG CTC AAC |
| D117H | GTT GAG CGG AGC CAG CTG CAC ATC TCA GAC CCG TAC AAA GTG TAC AG | CT GTA CAC TTT GTA CGG GTC TGA GAT GTG CAG CTG GCT CCG CTC AAC |
| K123R | CTG GAC ATC TCA GAC CCG TAC AGA GTG TAC AGG ATT GTT CCT GAG G | C CTC AGG AAC AAT CCT GTA CAC TCT GTA CGG GTC TGA GAT GTC CAG |

**Note S1:**

Tag sequences:
His8-linker-His8:
MGHHHHHHHHGATGSTAGSGTAGSTGASGASTGGTGATHHHHHHHHENLYFQGGS
GST-3C:
MSPILGYWKIKGLVQPTRLLLEYLEEKYEEHLYERDEGDKWRNKKFELGLEFPNLPYYIDGD
VKLTQSMAIIRYIADKHNMLGGCPKERAEISMLEGAVLDIRYGVSRIAYSKDFETLKVDFLSK
LPEMLKMFEDRLCHKTYLNGDHVTHPDFMLYDALDVVLYMDPMCLDAFPKLVCFKKRIEAI
PQIDKYLKSSKYIAWPLQGWQATFGGGDHPPKSDLEVLFQGPMGHM

**Figure S1**

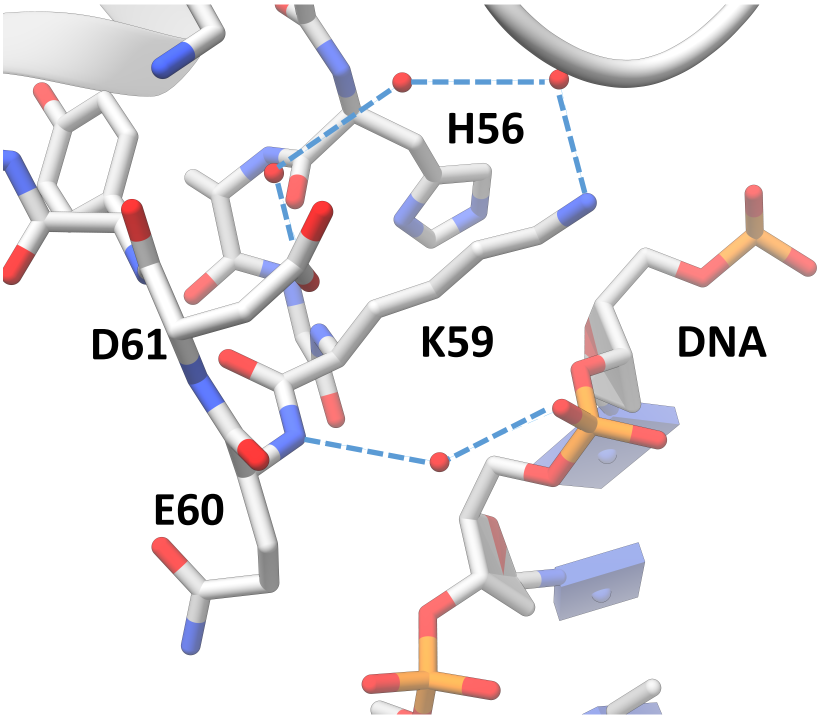


**Figure S1**: K59 lies on the surface of IRF4, close to the DNA-binding interface. Mutation to R or N may alter DNA binding.

**Figure S2**

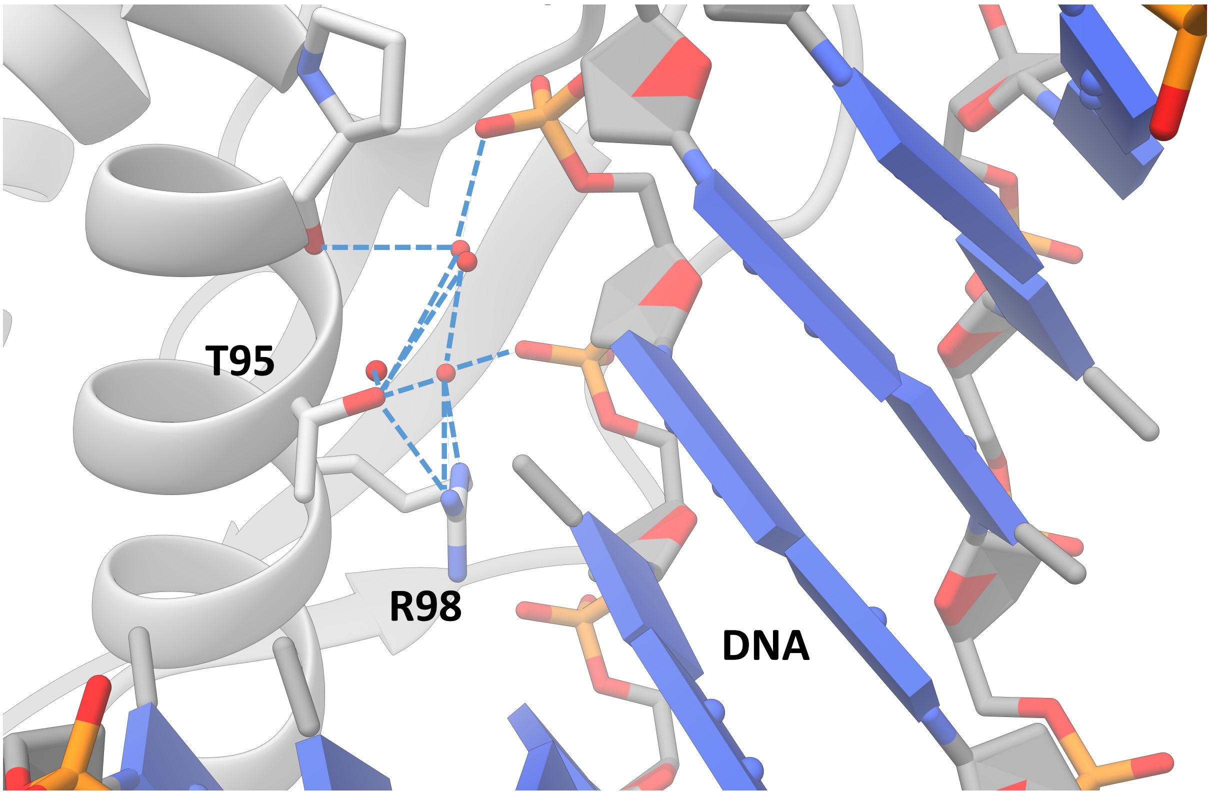


**Figure S2:** T95 lies on the surface of the IRF4 DNA-binding domain, close to the DNA-binding interface. The side-chain hydroxyl of T95 is involved in a complex network of hydrogen bonds, including water-mediated H-bonds to the DNA phosphate backbone. Mutation to A, K or R would be predicted to alter interaction with DNA.

**Figure S3**

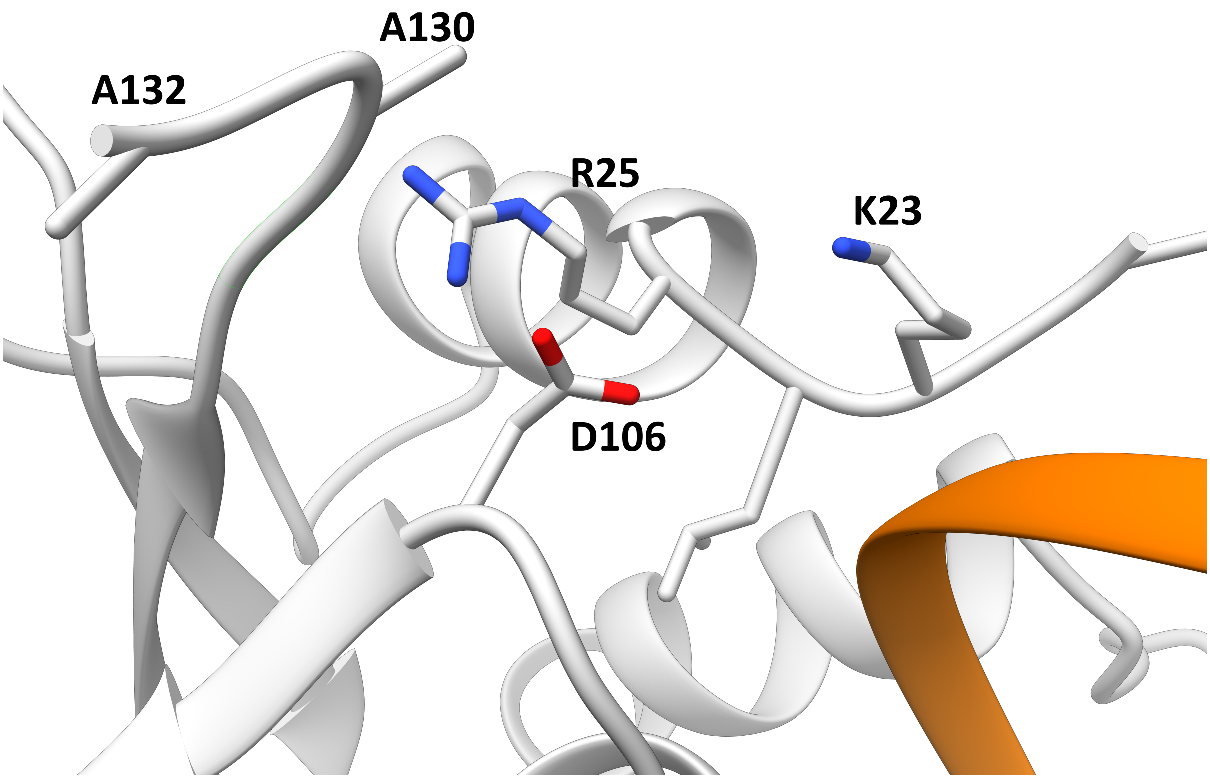


**Figure S3:** D106 lies close to the surface of the IRF4 DNA binding domain in proximity to the DNA binding interface and the N- and C-termini. Mutation to tyrosine could affect DNA binding positively or negatively**.**

**Figure S4**

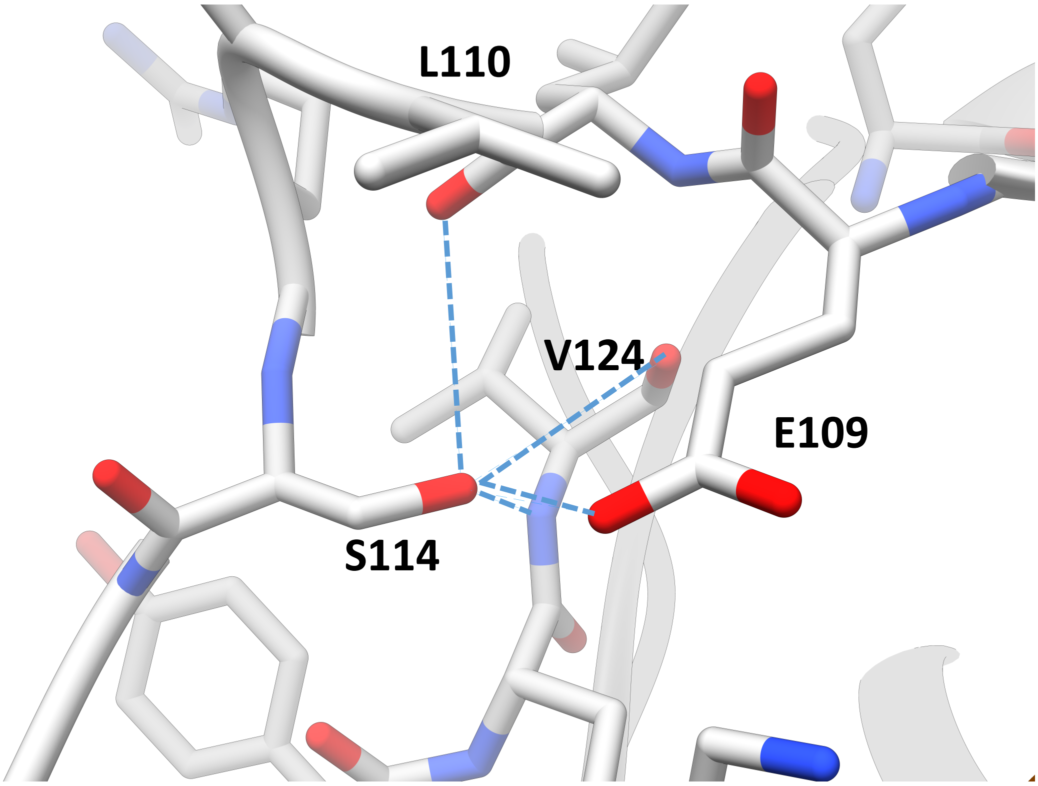


**Figure S4:** S114 is involved in a network of H-bonds with neighbouring residues and lies close to the DNA-binding interface. Mutation to R or N would disrupt this network of H-bonds and destabilise the DNA-binding interface.

**Figure S5**

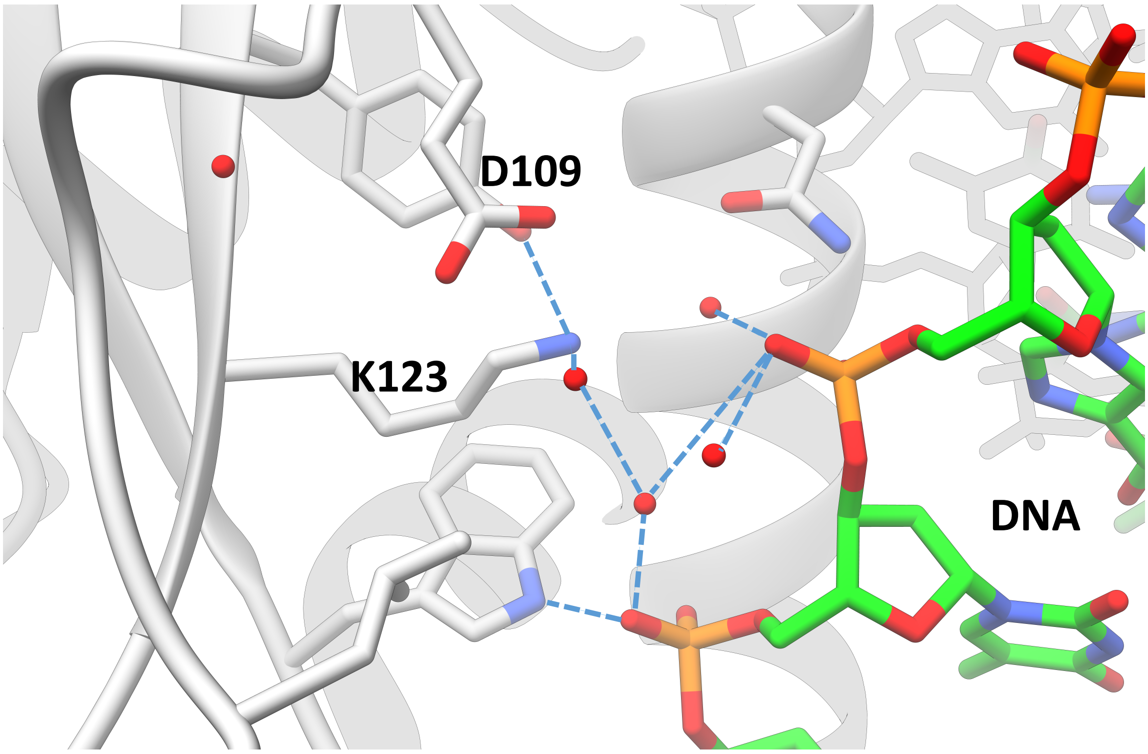


**Figure S5**: K123 lies close to the DNA binding interface. The side-chain amino of K123 is involved in a network of H-bonds, including water-mediated H-bonds to the DNA phosphate backbone. Mutation to R would be predicted to alter DNA binding.

**Figure S6**

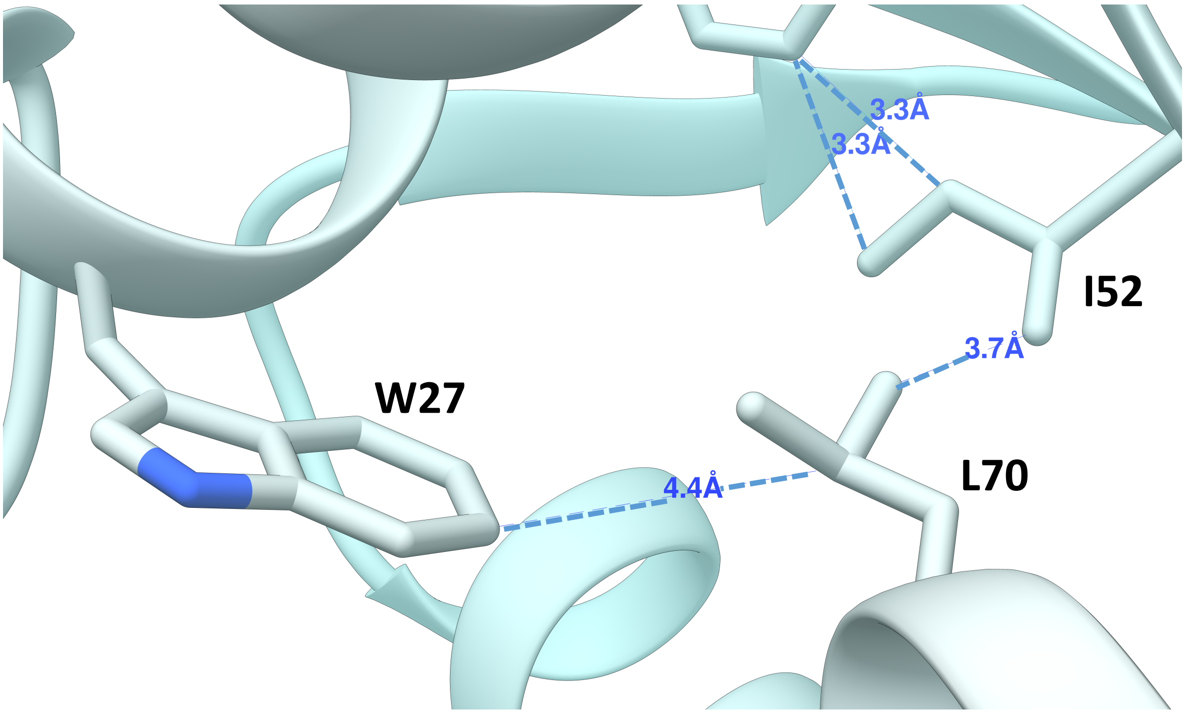


**Figure S6**: L70 lies within the IRF4 DNA-binding domain hydrophobic core. Mutation to valine is likely to reduce the stability of the core fold through loss of contacts with W27 and I52.

**Figure S7**

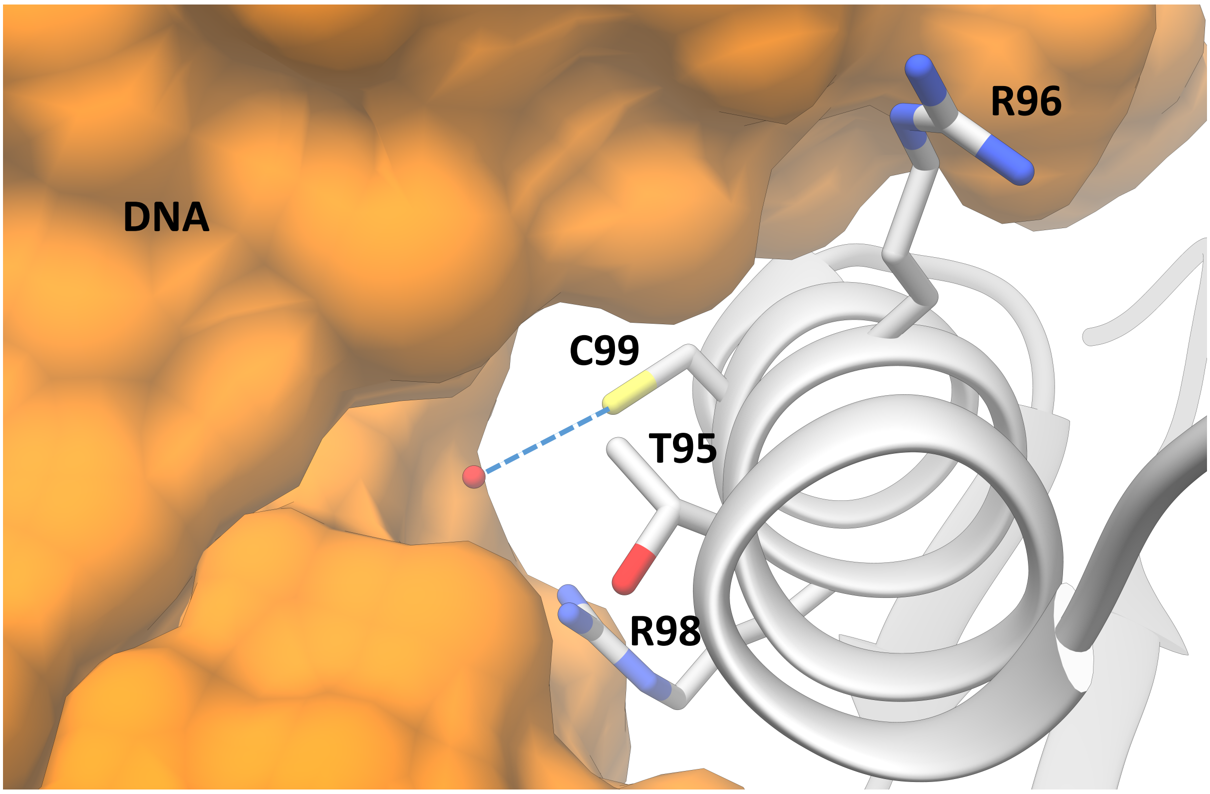


**Figure S7:** C99 lies on the surface of the IRF4 DNA-binding domain, within the DNA-binding interface. Mutation to R would be predicted to disrupt interaction with DNA; mutation to S should be tolerated.

Figure S8

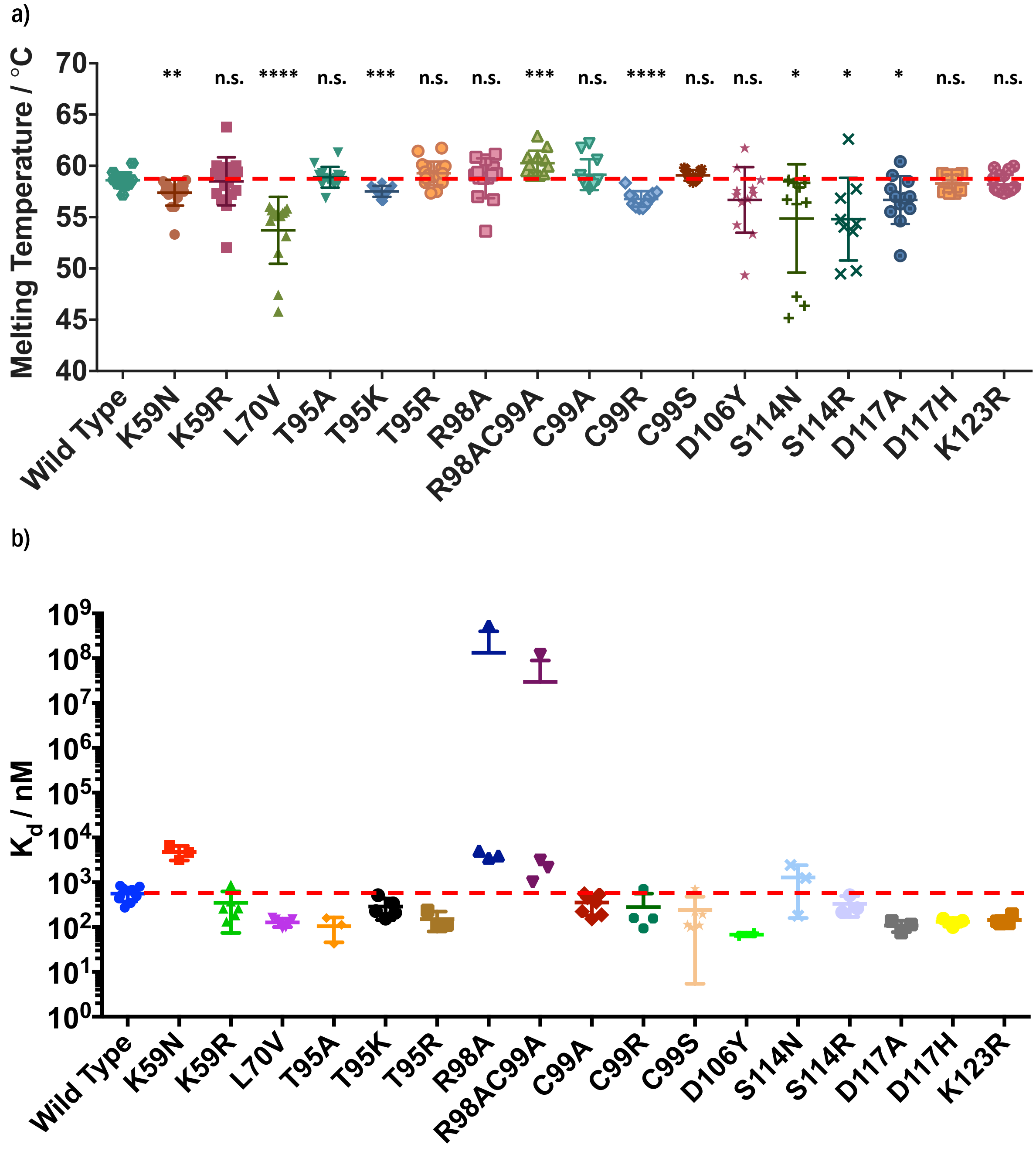


**Figure S8.** (**a**) Spread of melting temperatures for each mutant shown as scatter plot with mean and standard deviation. Dashed red line represents mean value for wild-type. (**b**) Results of fluorescence polarisation assay for WT and mutant IRF4 binding to IRE shown as scatter plot with mean and standard deviation. Dashed red line represents mean value for wild-type. Exemplar binding curves presented as supplementary figure S10.

**Figure S9**

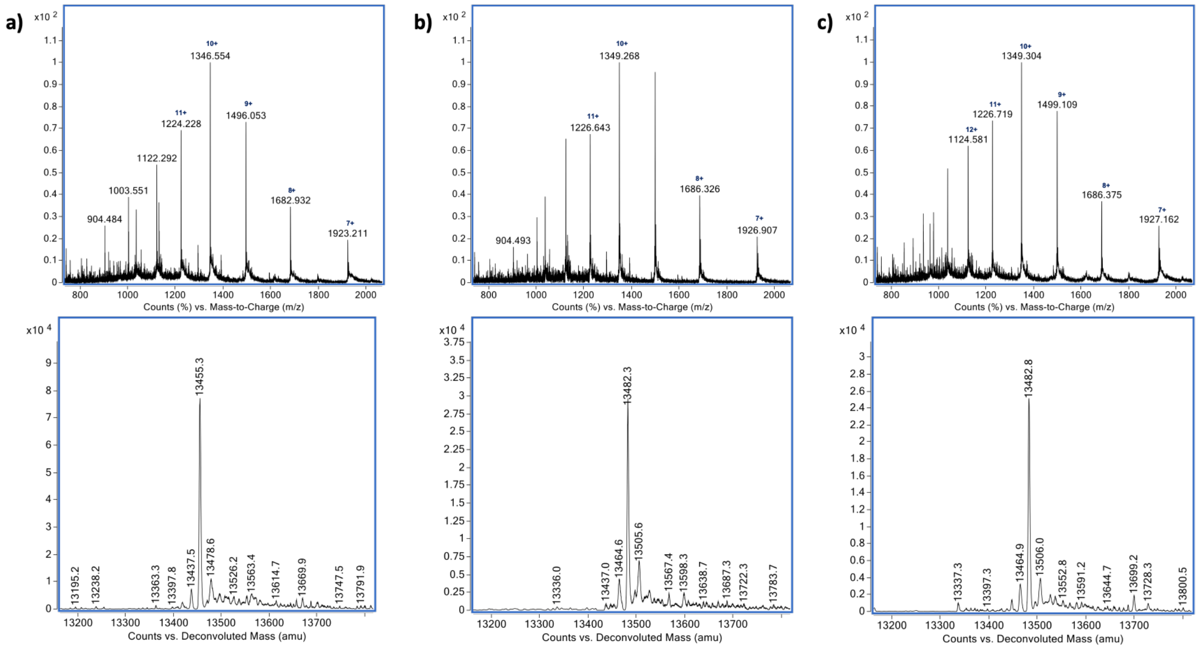


**Figure S9**: LC-MS analysis of intact IRF4 proteins. ESI-QTOF raw and deconvoluted mass spectra for (a) wild-type, (b) K59R and (c) K123R

**Figure S10**

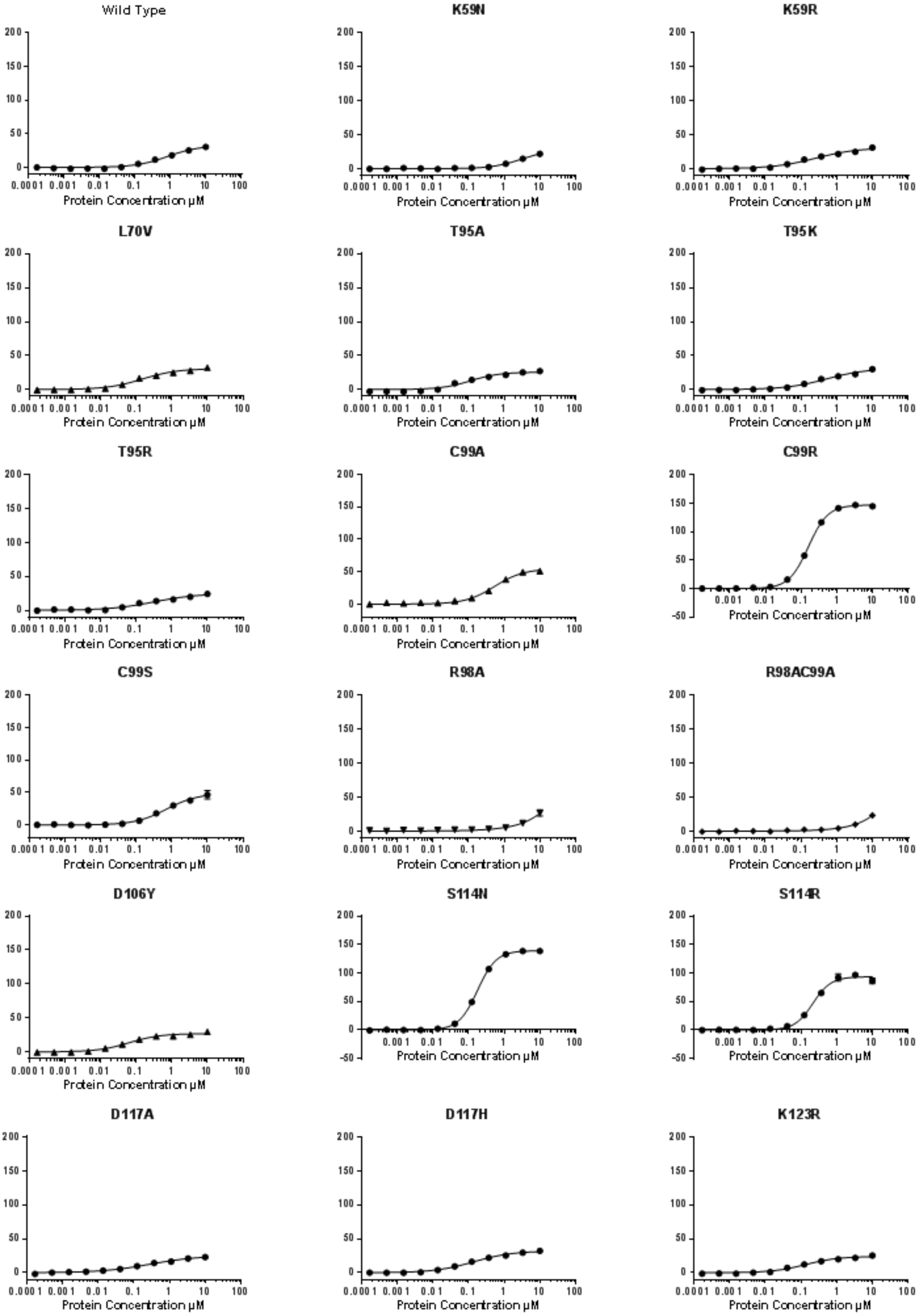


**Figure S10**: Exemplar fluorescence polarisation curves from two independent batches of protein, referenced against a non-protein control containing FAM-labelled IRE hairpin in assay buffer.

**Figure S11**

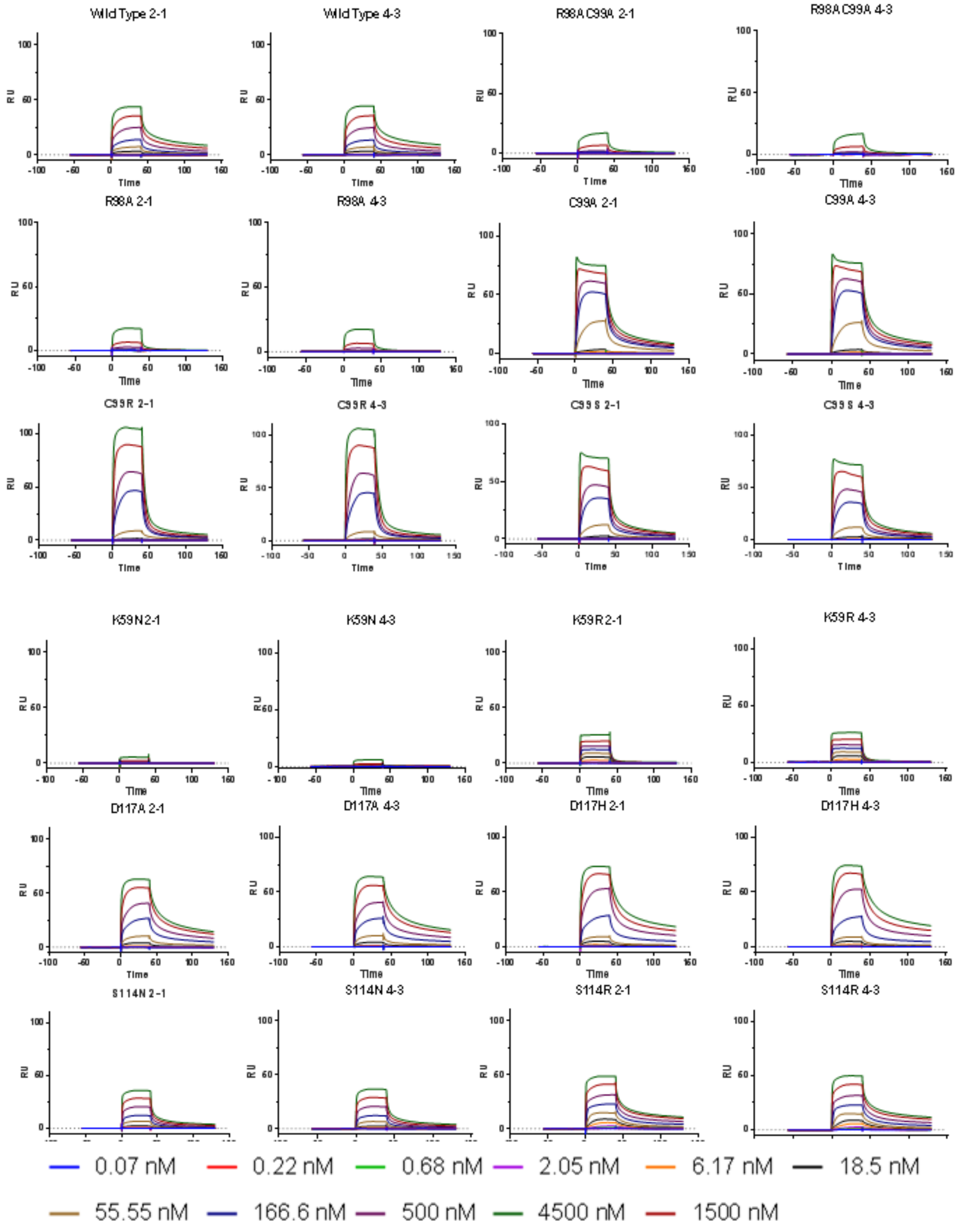


**Figure S11**: Exemplar SPR sensorgrams for wild type, R98, C99, R98A/C99A, K59, D117 and S114 mutants referenced to unliganded streptavidin (2-1) or scrambled hairpin, scrIRE (4-3) surfaces.

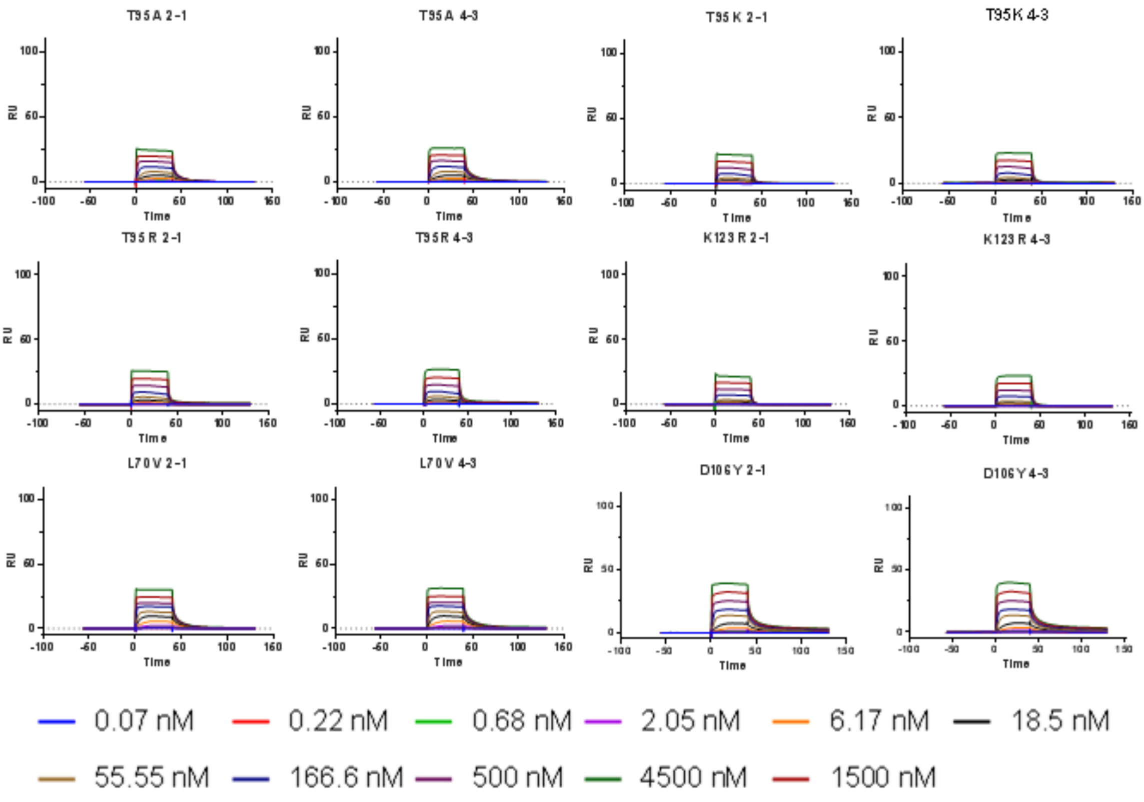


**Figure S11 cont**.: T95, K123, L70 and D106 mutant sensorgrams, referenced to streptavidin (2-1) and scrIRE (4-3).

**Figure S12**

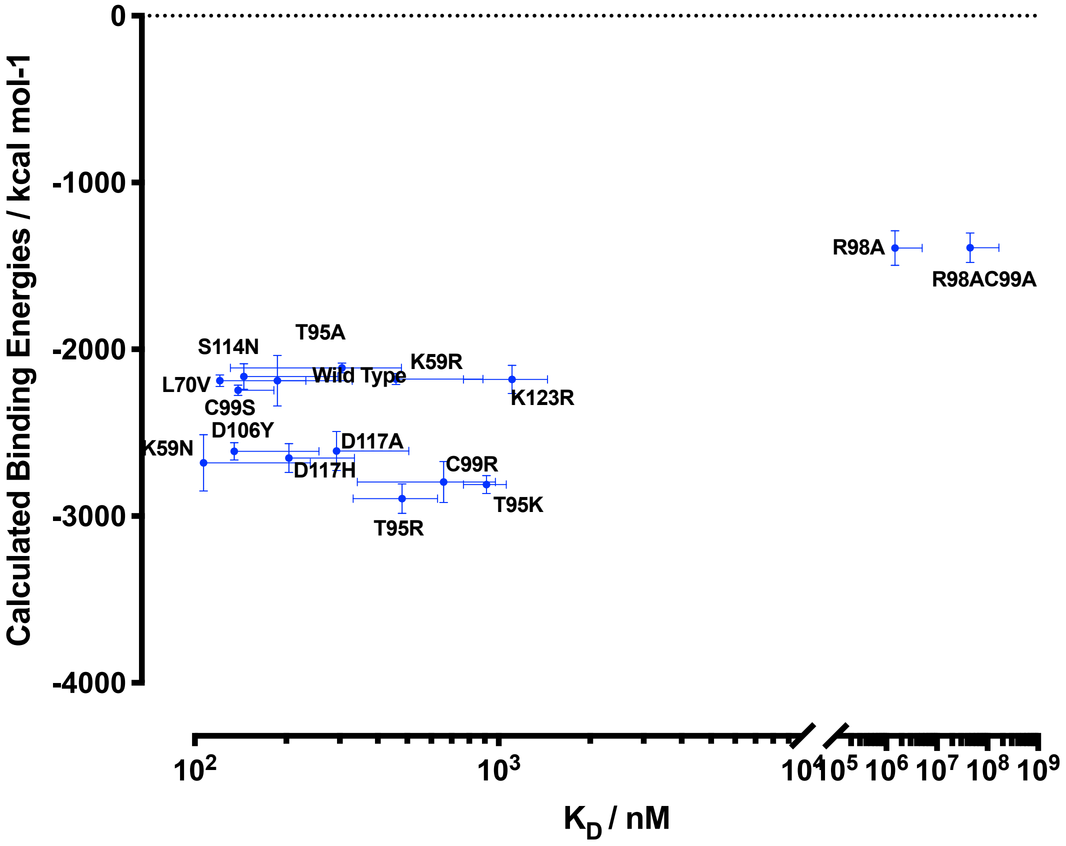


**Figure S12**: Comparison of computationally-derived (MM/PBSA) binding energy with experimental affinity (by SPR, *kd1*, two-state binding affinity).

### Figure S13

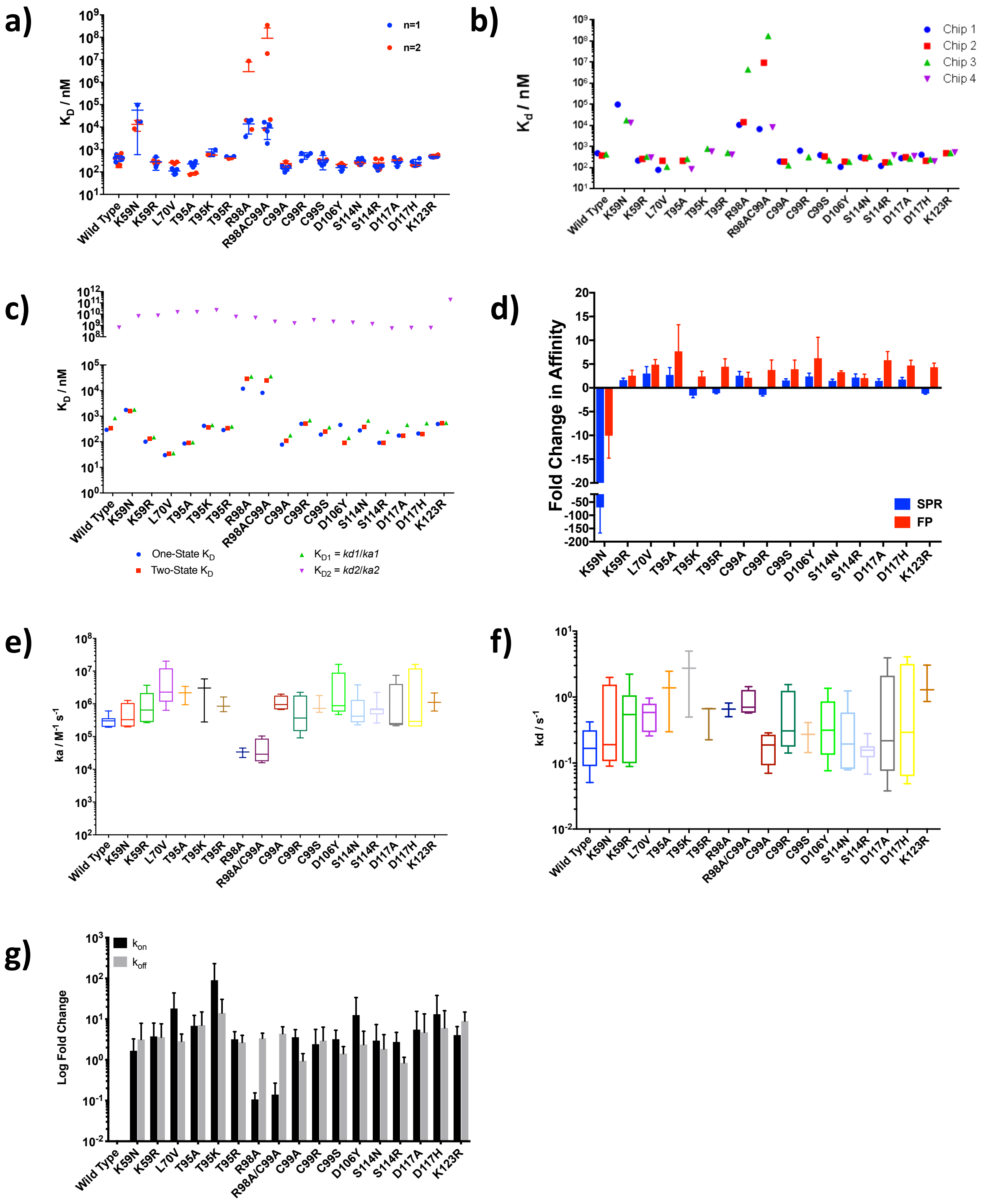


**Figure S13:** (**a**) Experimental affinity as determined by SPR, coloured for biological replicate variation, shown as scatter plot depicting mean and standard variation for each batch per protein. (**b**) Experimental affinity as determined by SPR, coloured to indicate variation between chip surfaces, shown as a scatter plot. (**c**) Comparison of one-state and two-state kinetic fitting to SPR data, demonstrating a weak second binding event contribution to overall $K_D$. Exemplar sensorgrams presented in figure S11. (**d**) Comparison of fold change in affinity to WT as measured by SPR (blue) and FP assay (red). (**e**) Association rate of each disease-associated mutant as determined by fitting two-state kinetic model to binding sensorgrams; (**f**) dissociation rate presented as box and whisker plot of mean and standard deviation; **(g)** log fold change in association and dissociation rates in comparison to wild type rates. Exemplar sensorgrams shown in supplementary figure S11.

**Figure S14**

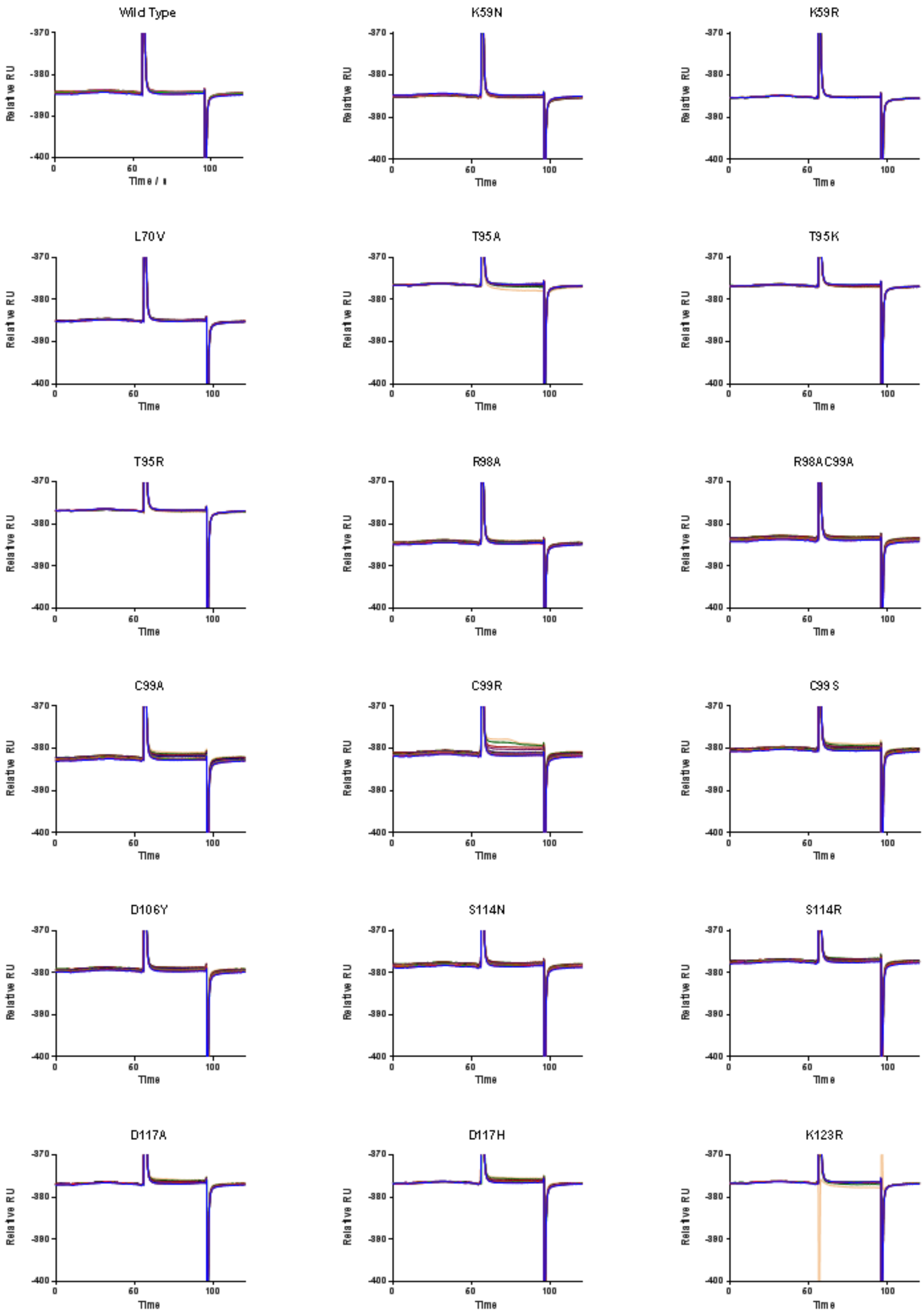

Wild Type
K59N
K59R
L70V
T95A
T95K
T95R
R98A
R98AC99A
C99A
C99R
C99S
D106Y
S114N
S114R
D117A
D117H
K123R
Relative RU
Time
-370
-380
-390
-400
0
60
100

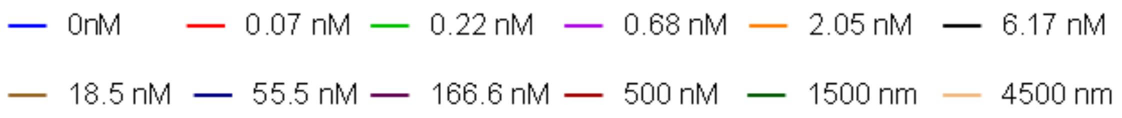


**Figure S14:** Sensorgram curves (3-1) for all variants and wild-type, demonstrating non-specific binding. Signal from channel with immobilised scrambled DNA (scrIRE) is referenced to streptavidin-only surface.

**Figure S15**

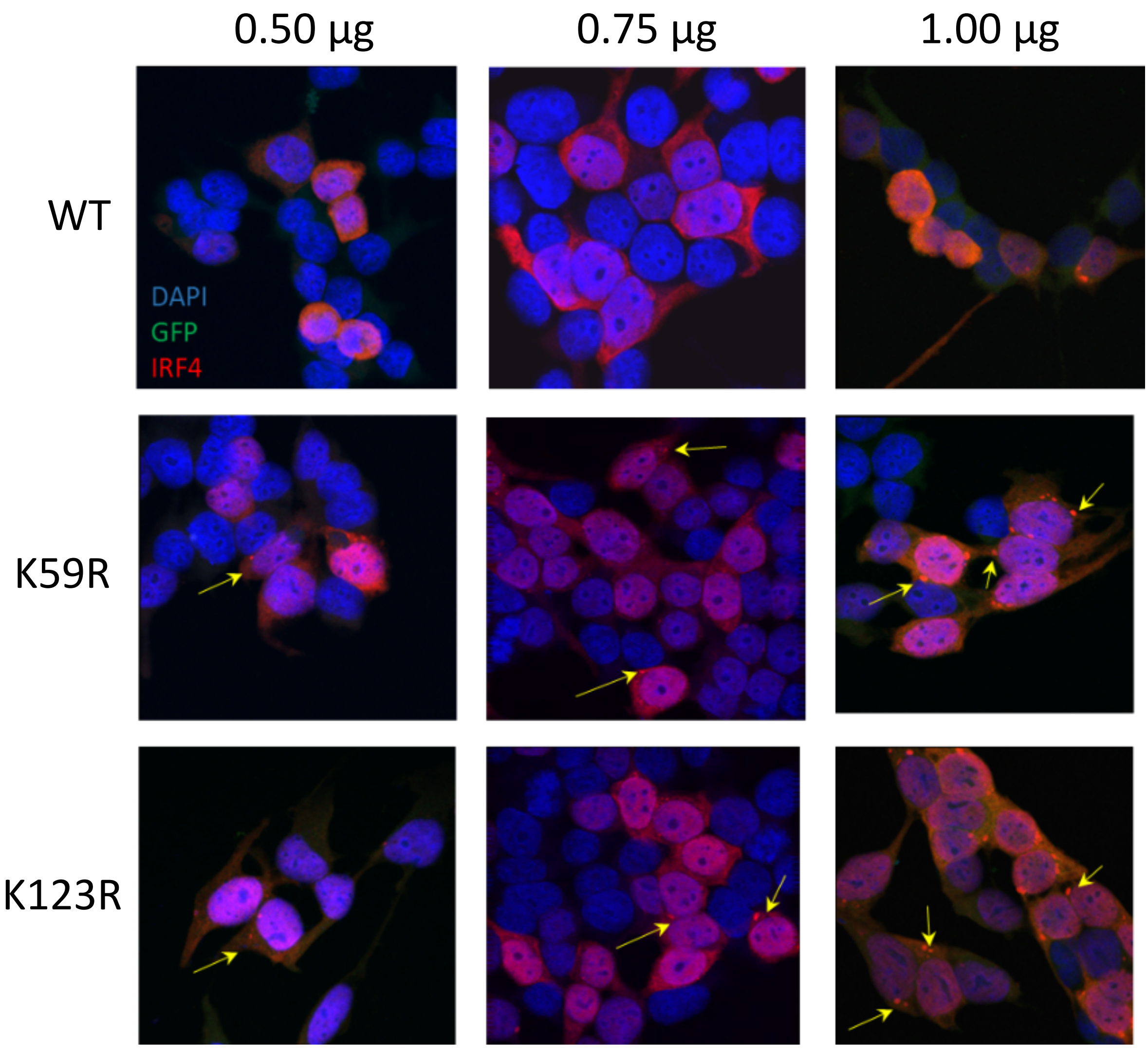


**Figure S15:** Immunofluorescence demonstrates differential sub-cellular localisation for IRF4 WT (upper panels), K59R (central panels) and K123R (lower panels). Nuclear staining (DAPI, blue), cytoplasmic staining (GFP, green) and IRF4 (red) and an image overlay (merge) at each concentration of plasmid is shown. Arrows depict cytoplasmic aggregates.
